# Phosphorylated transcription factor PuHB40 is involved in ROS-dependent anthocyanin biosynthesis in pear exposed to high-light stress

**DOI:** 10.1101/2023.12.22.573105

**Authors:** Lu Zhang, Lu Wang, Yongchen Fang, Yuhao Gao, Shulin Yang, Jun Su, Junbei Ni, Yuanwen Teng, Songling Bai

## Abstract

As sessile organisms, plants are increasingly vulnerable to environmental stresses because of global warming and climate change. Stress-induced reactive oxygen species (ROS) accumulation results in plant cell damages and even cell death. Anthocyanins are important antioxidants that scavenge ROS to maintain ROS homeostasis. However, the mechanism underlying ROS-induced anthocyanin accumulation is unclear. In this study, we determined that in pear the HD-Zip I family member PuHB40 mediates ROS-dependent anthocyanin biosynthesis under high-light stress. Specifically, PuHB40 is a transcription factor that induces PuMYB123-like/PubHLH3 complex for anthocyanin biosynthesis. The transcriptional activation by PuHB40 depends on its phosphorylation level, which is regulated by protein phosphatase 2A (PP2A). High ROS contents maintain the phosphorylation of PuHB40 at a high level, while also enhancing PuHB40-induced *PuMYB123-like* transcription by decreasing the transcription of *PuPP2AA2*, ultimately leading to increased anthocyanin biosynthesis. Our study revealed the pathway regulating ROS-induced anthocyanin biosynthesis in pear, further clarifying the mechanism underlying abiotic stress-induced anthocyanin biosynthesis, which may have implications for improving plant stress tolerance.

**IN A NUTSHELL:** *Background:* Various abiotic stresses, including high-light intensity, promote the accumulation of anthocyanins in plants, while also activating the production of reactive oxygen species (ROS). Anthocyanins can attenuate the negative effects of high-light stress by acting as antioxidants that restrict ROS accumulation. Several reports have shown that ROS can stimulate anthocyanin accumulation, but whether high-light stress-induced anthocyanin accumulation depends on ROS is undetermined. Additionally, the mechanism underlying ROS-dependent anthocyanin biosynthesis remains unclear.

*Question:* Does high-light stress-induced anthocyanin biosynthesis depend on ROS? What is the molecular basis of high-light stress-induced anthocyanin biosynthesis?

*Findings:* High-light stress-induced anthocyanin biosynthesis in pear seedlings is dependent on ROS accumulation. PuMYB123-like is the key MYB transcription factor for anthocyanin biosynthesis, while PuHB40 activates anthocyanin biosynthesis in response to ROS under high-light stress. Specifically, PuHB40 activates the transcription of *PuMYB123-like*, with the encoded protein combining with PubHLH3 to form an MBW complex that promotes anthocyanin biosynthesis. The transcriptional activation by PuHB40 depends on its phosphorylation status, which is regulated by protein phosphatase 2A (PP2A). High ROS levels inhibit the transcription of *PuPP2AA2*, thereby maintaining the phosphorylation of PuHB40, enhancing the transcriptional activation by PuHB40, inducing *PuMYB123-like* transcription, and ultimately leading to increased anthocyanin biosynthesis.

*Next steps:* Our future research will focus on whether ROS and the MYB123-like– PuHB40–PP2A regulatory module are also involved in the anthocyanin biosynthesis induced by other abiotic and biotic stresses, which will provide insights into biotic stress-induced anthocyanin biosynthesis and form the theoretical basis for improving fruit coloration.

## INTRODUCTION

Because of global warming and climate change, plants are increasingly becoming more susceptible to a variety of environmental stresses (e.g., drought, salinity, cold, and high-light) during growth and development, which leads to changes in physiological and biochemical metabolism, including the dramatic accumulation of reactive oxygen species (ROS) (Hipsch et al. 2021; Mittler et al. 2022). Light is one of the most indispensable environmental factors for plants. It provides energy for photosynthesis, which converts carbon dioxide (CO2) and water into energy-rich organic matter required for plant physiological metabolism (Croce and van Amerongen 2014). Under high-light stress conditions, photon flux exceeds the energy needed by plants to fix CO2, with O2^−^ and 1O^2^ primarily produced in chloroplasts by photosystems I and II, respectively, while H2O2 is generated in peroxisomes if photorespiration is activated (Galvez-Valdivieso et al. 2009; Kerchev et al. 2016; Wang et al. 2020; Mittler et al. 2022). Excessive ROS accumulation adversely affects the surrounding biological system components, including membranes, proteins, and DNA, and then triggers physiological processes that lead to programmed cell death (Lee and Gould 2002; Mittler 2017). Anthocyanins are important specialized metabolites that can function as non-enzymatic antioxidants that maintain ROS homeostasis and limit the damage due to stress-induced ROS. Moreover, their antioxidant capacity is greater than that of other flavonoids or even ascorbic acid and vitamin E (Gould et al. 2002; Bi et al. 2014; Acero et al. 2019). Considering various abiotic stresses promote anthocyanin accumulation in plants (Catalá et al. 2011; Xie et al. 2012; Jaakola 2013; Nakabayashi et al. 2014) and ROS can induce anthocyanin production (Xu et al. 2017; Shi et al. 2018), ROS may influence abiotic stress-induced anthocyanin biosynthesis; this hypothesis has been tested in a few studies (Wu et al. 2016; Qu et al. 2018; Shi et al. 2022).

Anthocyanins are synthesized in the phenylpropanoid pathway by various enzymes encoded by structural genes. Examples include phenylalanine ammonia-lyase (PAL), chalcone synthase (CHS), chalcone isomerase (CHI), flavanone 3-hydroxylase (F3H), flavonoid 3′-hydroxylase (F3′H), dihydroflavonol 4-reductase (DFR), anthocyanin synthase (ANS), and UDP-glucose: flavonoid 3-glucosyltransferase (UFGT) (Xu et al. 2015; Liu et al. 2021; Pratyusha and Sarada 2022). The R2R3-MYB and basic helix–loop– helix (bHLH) transcription factors (TFs) combine with WD-repeat proteins to form MBW complexes in which MYBs function as major regulators of structural gene expression (Jaakola 2013; Xu et al. 2015; Naing and Kim 2018). The R2R3-MYB TFs are divided into 25 subgroups, among which subgroup 6 contains TFs that are mainly responsible for promoting anthocyanin biosynthesis, including PAP1 (AtMYB75), PAP2 (AtMYB90), and AtMYB113 in *Arabidopsis thaliana* (Dubos et al. 2010) as well as PpMYB10 and PpMYB114 in pear (Feng et al. 2010; Yao et al. 2017). The expression of MYB TFs is regulated by various factors, including light (Takos et al. 2006; Li et al. 2012; Bai et al. 2019a), low/high temperatures (Choi et al. 2009; Lin-Wang et al. 2011; Xie et al. 2012), and phytohormones (Shan et al. 2009; Das et al. 2012; Ni et al. 2023). The encoded MYB TFs control the expression of anthocyanin biosynthetic structural genes. To date, the molecular basis of ROS-induced anthocyanin biosynthesis remains relatively uncharacterized.

In cells, the accumulation of ROS alters the redox state and the post-translational modification of proteins, especially the phosphorylation of proteins. ROS signal is transduced by Ca^2+^, Ca^2+^ binding protein, phosphatidic acid (PA) and phospholipase D (PLD) and activates OXIDATIVE SIGNAL-INDUCIBLE 1 (OXI1), which subsequently activates MAPK cascade pathway, which in turn promotes the phosphorylation of downstream transcription factors, such as WRKY proteins and AP2/ethylene-response factors (ERFs) (Kovtun et al. 2000; Jalmi and Sinha 2015; Byrne et al. 2020; Mittler et al. 2022). Furthermore, the phosphorylation of transcription factors are also regulated by phosphatase, such as PTP1 (protein tyrosine phosphatase 1), SAL1 (3’-phosphoadenosine 5’-phosphate phosphatase) and PP2C, whose activity are regulated by ROS levels (Gupta and Luan 2003), oxidative stresses (Chan et al. 2016) or phytohormones (Mine et al. 2017). The PP2A holoenzyme, which is a serine/threonine-specific phosphatase, is a heterotrimeric complex comprising a catalytic subunit C, a regulatory subunit B, and a structural subunit A. The A subunit has a conserved structure consisting of 15 tandem repeats of a 39 amino acid sequence (i.e., HEAT motif) (Janssens and Goris 2001). In *A. thaliana*, three A subunits of PP2A have been identified, namely PP2AA1 (RCN1), PP2AA2, and PP2AA3. The phosphatase activity of PP2A plays an important role in stress response (DeLong 2006; Uhrig et al. 2013; Lillo et al. 2014; Durian et al. 2016). However, it is not clear whether PP2A phosphatases are involved in ROS regulation of protein phosphorylation under stress conditions.

In this study, we determined that abiotic stress-induced anthocyanin biosynthesis is at least partially dependent on ROS in pear. The HD-Zip I family member PuHB40 was identified as the key TF involved in ROS-dependent anthocyanin biosynthesis. Notably, PuHB40 functions as the upstream positive regulator of the PuMYB123-like/PubHLH3 complex during anthocyanin biosynthesis. The dephosphorylation of PuHB40, which is catalyzed by protein phosphatase 2A (PP2A), inhibits its function related to ROS-induced anthocyanin biosynthesis. Under abiotic stress conditions, ROS inhibit the transcription of *PuPP2AA2*. This study revealed the pathway regulating ROS-induced anthocyanin biosynthesis in pear under high-light stress, providing important insights into the mechanism mediating abiotic stress-induced anthocyanin production.

## RESULTS

### Stress induced ROS promoted anthocyanin biosynthesis in pear seedlings

To test whether abiotic stress-induced anthocyanin biosynthesis depends on ROS production, pear seedlings (*P. ussuriensis*) grown under continuous light were exposed to abiotic stresses, including salt, drought, and high-light, and the N,N′-dimethylthiourea (DMTU) was used to decrease the ROS content under abiotic stress conditions. DMTU is a highly permeant molecule that can scavenge hydrogen peroxide (H2O2), the hydroxyl radical (·OH), or hypochlorous acid (HOCl) *in vitro* (Curtis et al. 1988; Serobatse and Kabanda 2017). It has been widely used to remove ROS in plant studies (Ni et al. 2019; Deng et al. 2022). The various abiotic stresses caused anthocyanins to accumulate in pear seedlings after seven days, but anthocyanin production decreased significantly following the DMTU treatment (Supplemental Fig. S1A), indicating that ROS are potentially involved in stress-induced anthocyanin production. Furthermore, ROS accumulation and anthocyanin biosynthesis were examined at 0, 6, 12, 24, 48, 72, and 96 h after plants underwent the high-light treatment (Fig. 1A). Anthocyanin accumulation peaked at the 72-h time-point and then decreased, but this accumulation was significantly suppressed in the seedlings that were also treated with DMTU (Fig. 1B). The NBT and DAB staining results indicated that DMTU suppressed the accumulation of O2^−^ and H2O2 in pear leaves exposed to high-light stress (Fig. 1C–F). These findings were supported by the analysis of H2O2 contents (Fig. 1G). Moreover, in response to the different stress treatments (e.g. high-light stress, drought stress, and salt stress), the anthocyanin and H2O2 contents accumulated in the same leaf regions (Supplemental Fig. S1B). According to these results, abiotic stress-induced anthocyanin accumulation depends on ROS production. Hence, the ROS scavenger DMTU can inhibit ROS accumulation and indirectly decrease anthocyanin production.

**Figure 1.**
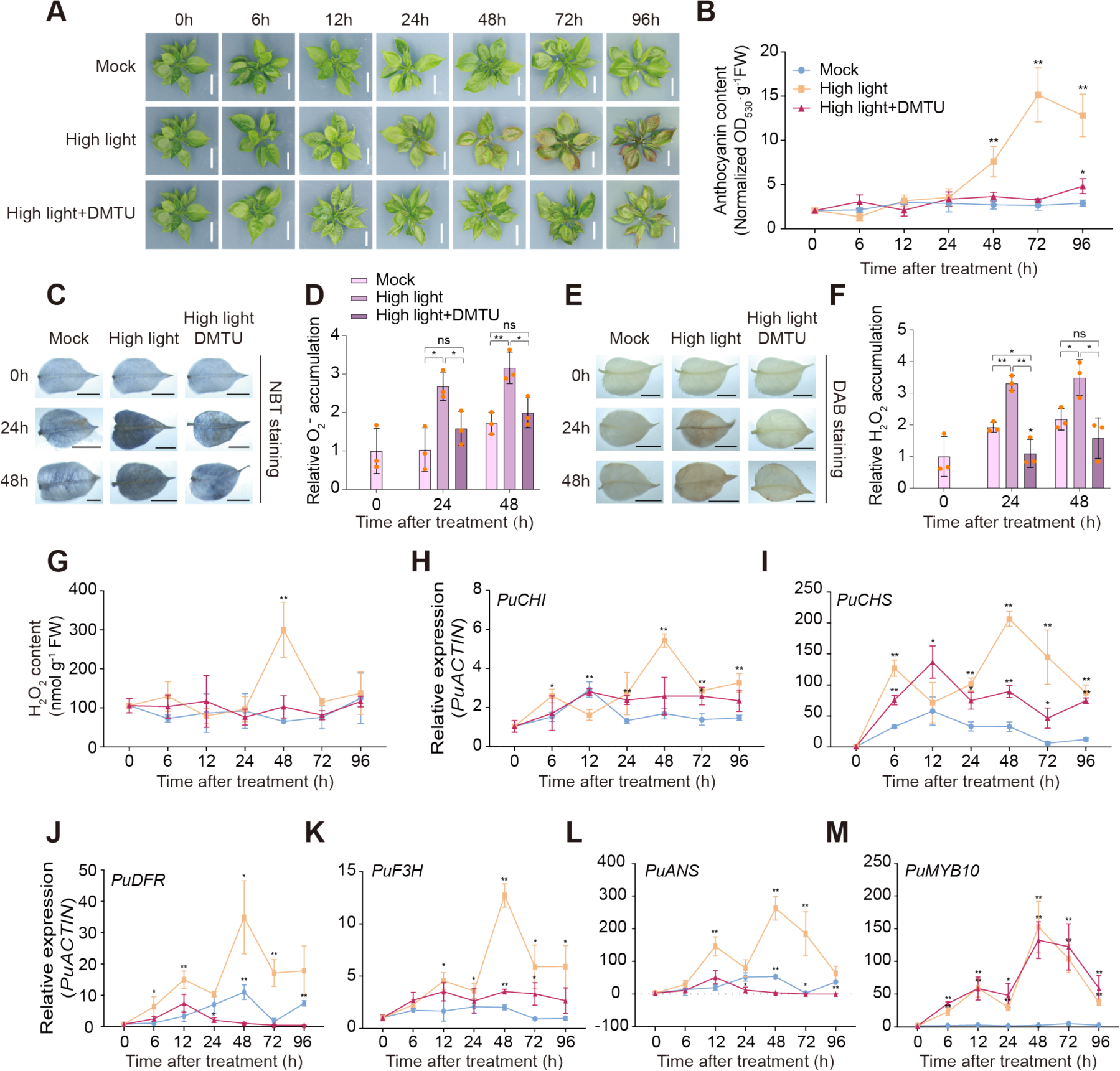
High-light stress-induced anthocyanin biosynthesis is ROS-dependent. **(A-M),** Pear seedlings were treated with high-light (photon flux density: approximately 160 μmol·s^-1^·m^-2^) or high-light supplemented with N, N’-dimethylthiourea (DMTU) treatment. DMTU was applied as ROS scavenger, and pear seedlings under normal light condition were used as Mock (photon flux density: approximately 50 μmol·s^-1^·m^-2^). Pear seedlings were sampled at 0 h, 6 h, 12 h, 24 h, 48 h, 72 h, 96 h. **(A)** Phenotype of pear seedlings (scale bar, 1 cm). **(B)** The anthocyanin contents for pear seedlings (units: A530/g of fresh weight). **(C)** NBT staining for O2^-^ in pear seedlings leaves at 0 h, 24 h, 48 h (scale bar, 5 mm). **(D)** Quantification of NBT staining in pear leaves at 0 h, 24 h, 48 h. **(E)** DAB staining for H2O2 in pear seedlings leaves at 0 h, 24 h, 48 h (scale bar, 5 mm). **(F)** Quantification of DAB staining in pear leaves at 0 h, 24 h, 48 h. **(G)** The H2O2 content for pear seedlings. **(H-M),** The expression levels of *PuCHI*, *PuCHS*, *PuDFR*, *PuF3H*, *PuANS*, *PuMYB10* in pear seedlings under different treatments. *PuACTIN* was amplified as an internal control. Data are presented as means ± s.d. of 3 biological replicates. Asterisks indicate significant differences compared with Mock group **(B)** and **(G-M)**, Mock and High light stress **(D)** and **(F)** (two-tailed Student’s *t-test*, \**P* < 0.05, \*\**P* < 0.01; ns, no significance, *P* > 0.05), all *P* values are shown in Supplemental Data Set S7.

To identify the key genes responsible for abiotic stress-induced anthocyanin biosynthesis, the expression of anthocyanin-related structural and regulatory genes was analyzed. The *PuCHI*, *PuCHS*, *PuDFR*, *PuF3H*, and *PuANS* expression levels increased significantly and peaked at 48 h after the high-light treatment, which was in accordance with the changes in anthocyanin accumulation. The expression of these genes was significantly inhibited by DMTU (Fig. 1, H-L). In addition, the expression levels of high-light-responsive genes (*PuELIP1* and *PuELIP2*) were significantly upregulated following the high-light treatment with and without DMTU, indicative of the effects of high-light stress (Supplemental Fig. S1C and D). Interestingly, *PuMYB10*, which encodes a critical TF that regulates light-induced anthocyanin biosynthesis in pears, was similarly expressed following the high-light stress and DMTU treatments (Fig. 1M), even though these two treatments resulted in significant differences in anthocyanin accumulation and the expression of downstream structural genes. These findings suggest that other TFs may be involved in regulating ROS-dependent anthocyanin biosynthesis.

### Identification of PuMYB123-like and its interaction with PubHLH3

To determine which TFs participate in ROS-dependent anthocyanin biosynthesis, the high-light stress- and DMTU-treated pear seedlings underwent an RNA-seq analysis. A weighted gene co-expression network analysis was conducted using the fragments per kilobase of transcript per million reads mapped (FPKM) values of anthocyanin-related structural genes (*PuCHS*, *PuCHI*, *PuF3H*, *PuDFR*, and *PuANS*) as trait data for identifying genes that are co-expressed with anthocyanin biosynthetic genes. A total of 23 modules were identified, with the “purple” module most highly correlated with the trait data (Supplemental Fig. S2). The “purple” module included nine MYB TF genes, of which eight encoded R2R3-MYB TFs (Supplemental Data Set S5). Finally, *PuMYB123-like* (EVM0034726.1) was more significantly induced by high-light stress than the other seven R2R3-MYB TF genes (Supplemental Fig. S3). Accordingly, it was selected for further study. The *PuMYB123-like* expression level, which was consistent with that of anthocyanin biosynthesis structural genes, increased rapidly in response to high-light stress and peaked at 48 h, but the DMTU treatment inhibited *PuMYB123-like* expression (Fig. 2A). The multiple sequence alignment showed PuMYB123-like contains a typical R2R3 repeat in the N-terminal, with a bHLH motif in the R3 repeat region that can interact with bHLH TFs (Supplemental Fig. S4B). The subcellular localization analysis showed that the fluorescence of GFP-tagged PuMYB123-like was predominantly distributed in the nucleus, consistent with the characteristics of TFs (Fig. 2C). Considered together, these results indicate that PuMYB123-like is a transcriptional factor that may contribute to ROS-dependent anthocyanin biosynthesis.

**Figure 2.**
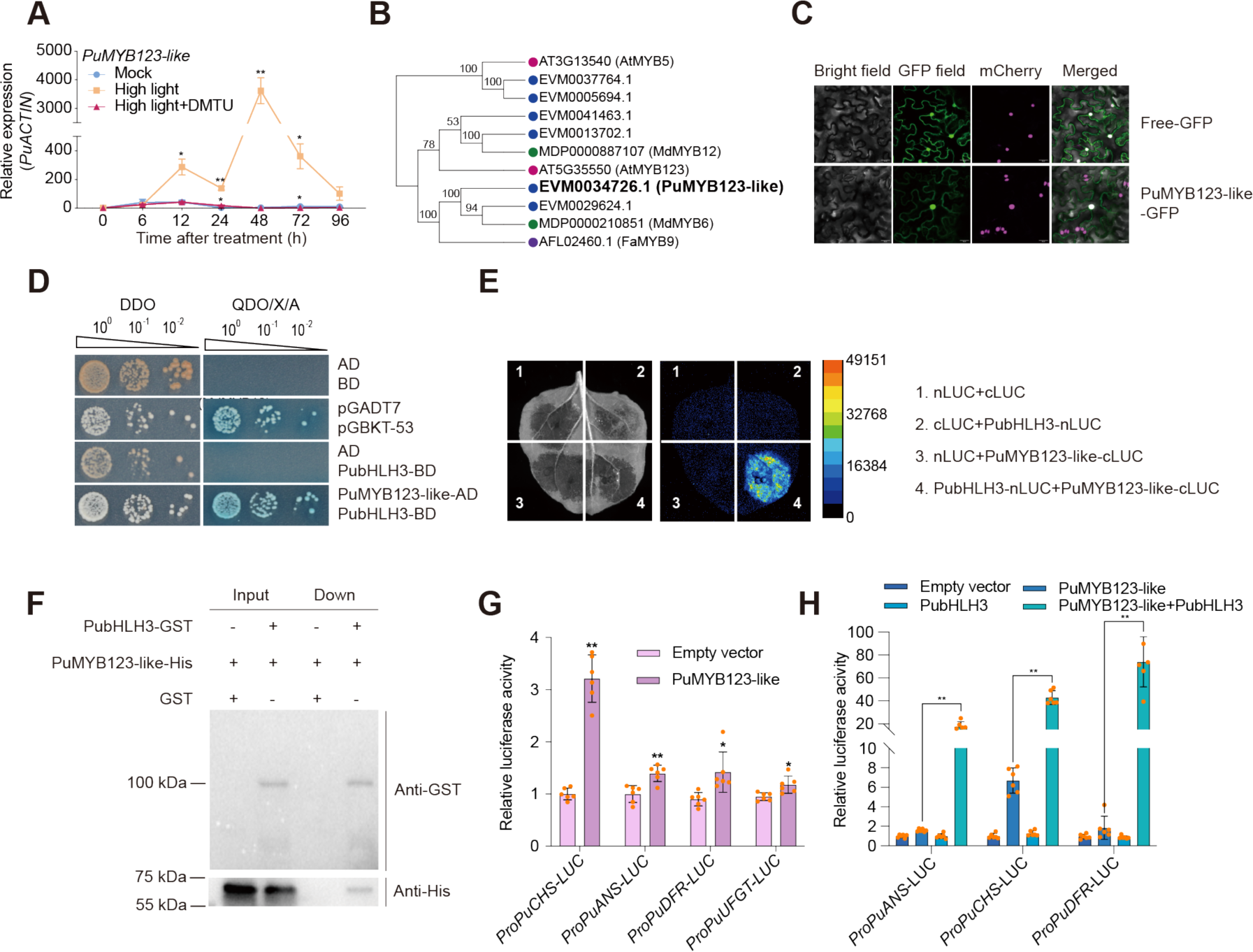
Identification of PuMYB123-like and its interaction with PubHLH3. **(A)** The expression levels of *PuMYB123-like* in pear seedlings under high-light treatment or high-light supplemented with DMTU treatment. The pear seedlings under normal light condition were used as Mock. *PuACTIN* was amplified as an internal control. **(B)** Phylogenetic analysis of PuMYB123-like. The phylogenetic tree was calculated using the maximum-likelihood method of MEGA X (version 10.1.8). Bootstrap values of 1000 replicates for each branch are shown. The protein sequences from pear (Pu, *Pyrus ussuriensis*, highlighted with blue dots), apple (Md, *Malus × domestica*, highlighted with green dots), Arabidopsis (At, *Arabidopsis thaliana*, highlighted with rose red dots) and strawberry (Fa, *Fragaria × ananassa*, highlighted with purple dots). PuMYB123-like (EVM0034726.1) is bolded and enlarged. **(C)** Subcellular localization of PuMYB123-like in *N. benthamiana* leaf cells (scale bar, 25 μm). **(D)** Interaction between PuMYB123-like and PubHLH3 in a yeast two-hybrid assay, with pGADT7-T and pGBKT7-53 as positive controls. DDO, SD/-Trp/-Leu medium; QDO/X/A, SD/-Trp/-Leu/-Ade/-His medium with X-α-gal and Aureobasidin. **(E)** The interaction between PuMYB123-like and PubHLH3 was detected by luciferase complementation imaging assays. The pseudocolour scale bar indicates the range of luminescence intensity. **(F)** The interaction between PuMYB123-like and PubHLH3 analyzed using a pull-down assay. PuMYB123-like-His was incubated with GST or PubHLH3-GST. The interaction proteins were precipitated by GST-Trap agarose beads and analyzed by immunoblotting with an anti-GST or anti-His antibody. **(G)** A dual-luciferase assay demonstrated that PuMYB123-like promotes the promoter activity of anthocyanin-related genes (*PuCHS*, *PuANS*, *PuDFR*, *PuUFGT*). The promoter of anthocyanin-related genes was cloned into the pGreenII 0800-LUC (firefly luciferase) vector, and the full-length CDS of *PuMYB123-like* was cloned into the pGreenII 0029 62-SK vector. The empty vector of pGreenII 0029 62-SK was used as control. **(H)** Dual-luciferase assay showed that interaction between PuMYB123-like and PubHLH3 significantly enhanced activation of PuMYB123-like on *PuCHS*, *PuDFR* and *PuANS* promoters. Data are presented as means ± s.d. of 3 biological replicates **(A)** or 6 biological replicates **(G)** and **(H)**. Asterisks indicate significant differences compared with Mock **(A)**, empty vector **(G)** or PuMYB123-like **(H)** (two-tailed Student’s *t-test*, \**P* < 0.05, \*\**P* < 0.01), all *P* values are shown in Supplemental Data Set S7.

Considering MYB and bHLH TFs combine with WD-repeat proteins to form MBW complexes that control the expression of anthocyanin-related structural genes, we assessed whether PuMYB123-like can interact with three candidate bHLH TFs (PubHLH3, PubHLH33, and PubHLH64), which affect anthocyanin biosynthesis in pear (Tao et al. 2020; Ni et al. 2021). According to the yeast two-hybrid (Y2H) assay results, PuMYB123-like can bind to PubHLH3 (Fig. 2D). The interaction was confirmed in pull-down and luciferase complementation imaging (LCI) assays, which revealed PuMYB123-like can interact with PubHLH3 both *in vivo* and *in vitro* (Fig. 2, E and F). Because of the self-activation of PubHLH33 and PubHLH64, the corresponding genes were fragmented to assess potential interactions. The results showed that PuMYB123-like can weakly interact with PubHLH33 (Supplemental Fig. S5A), and interact with the truncated PubHLH64 (encoded by 747–1,086 bp) (Supplemental Fig. S5B). According to the dual-luciferase assay, PuMYB123-like can significantly activate the promoter of the anthocyanin-related structural gene *PuCHS*, whereas it weakly activates the *PuANS*, *PuDFR*, and *PuUFGT* promoters (Fig. 2G). However, PubHLH3 significantly enhanced the activation of the *PuCHS*, *PuDFR*, and *PuANS* promoters by PuMYB123-like, whereas PubHLH33 and PubHLH64 had no significant effect (Fig. 2H; Supplemental Fig. S5C). Based on these observations, we proposed that PuMYB123-like, which belongs to subgroup 5 of the R2R3-MYB family, promotes anthocyanin-related structural genes expression as a component of the MBW complex that also includes PubHLH3.

### PuMYB123-like promotes anthocyanin biosynthesis in pears

To confirm PuMYB123-like affects anthocyanin biosynthesis, *PuMYB123-like* was overexpressed in pear calli. After a 3-day exposure to light, the anthocyanin contents and anthocyanin biosynthetic gene expression levels were significantly higher in the calli overexpressing *PuMYB123-like* than in the wild-type calli. In addition, the silencing of *PuMYB123-like* in pear calli via RNA interference (RNAi) inhibited anthocyanin biosynthesis and decreased the expression of anthocyanin biosynthetic genes (Fig. 3A-C). Considering PuMYB123-like belongs to subgroup 5 of the R2R3-MYB family, which reportedly regulates proanthocyanin biosynthesis (Baudry et al. 2004; Dubos et al. 2010), the proanthocyanin contents in *PuMYB123-like*-overexpressing (*PuMYB123-like*-OE) pear calli was also examined. There was no significant difference in the proanthocyanin contents of the wild-type and transgenic pear calli (Supplemental Fig. S6A). Additionally, *PuMYB123-like* was ectopically expressed in *Arabidopsis thaliana* to verify the function of the encoded TF. Compared with the wild-type control, the two independent transgenic lines accumulated more anthocyanin (Fig. 3D-F) and had shorter hypocotyls (Fig. 3G). The expression of the anthocyanin biosynthetic genes *AtCHS*, *AtCHI*, *AtDFR*, *AtF3H*, *AtLODX*, and *AtPAP1* increased significantly in the *PuMYB123-like*-expressing lines (Fig. 3H). To examine whether PuMYB123-like mediates anthocyanin biosynthesis in fruits, *PuMYB123-like* was transiently overexpressed in the peel of ‘Zaosu’ pear fruits that were bagged in advance. The comparison with the control peels (empty vector) after a 3-day exposure to light indicated the *PuMYB123-like*-OE peels had significantly higher anthocyanin contents (Supplemental Fig. S6, B and C), and the transcription level of anthocyanin biosynthetic gene (i.e., *PpCHS*, *PpDFR*, *PpF3H*, *PpANS*, *PpUFGT*, *PpMYB10*, and *PpMYB114*) (Supplemental Fig. S6D). Similarly, ‘Hongzaosu’ red pear fruit peels in which *PuMYB123-like* was transiently silenced via virus-induced gene silencing (VIGS) had lower anthocyanin contents and anthocyanin biosynthetic gene transcription levels than the control peels (TRV) (Supplemental Fig. S6, E-G). Considered together, these results demonstrate that PuMYB123-like positively regulates anthocyanin biosynthesis in pear.

**Figure 3.**
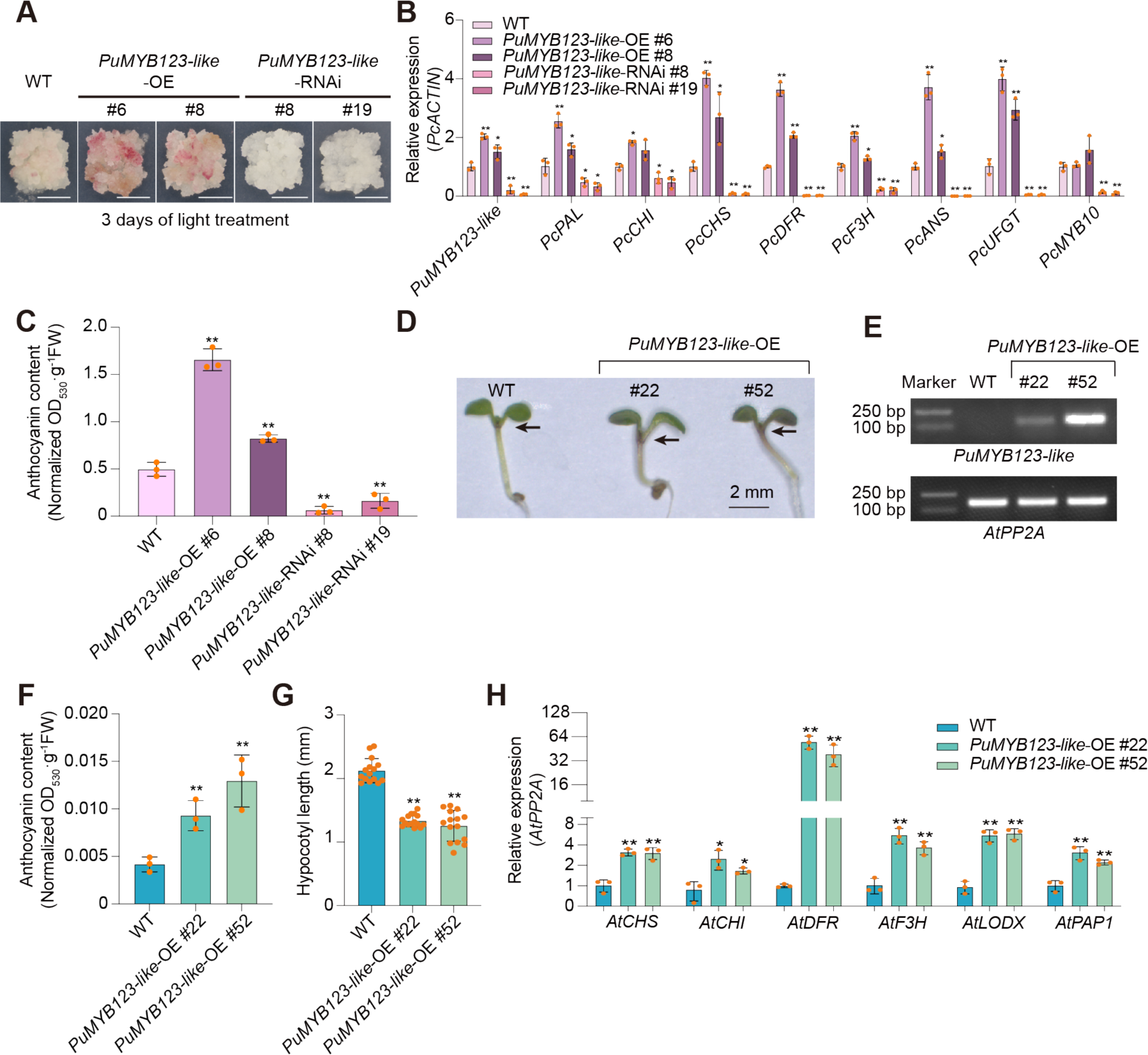
PuMYB123-like promotes anthocyanin biosynthesis. **(A)** Phenotype of pear calli overexpressing or silencing *PuMYB123-like* (scale bar, 1 cm). Wild type pear calli were used as the negative control. Pear calli were incubated under continuous white light at 17°C for 3 d. **(B)** Expression levels of anthocyanin-related genes and *PuMYB123-like* in *PuMYB123-like*-OE *and PuMYB123-like*-RNAi pear calli after light treatment. *PcACTIN* was amplified as an internal control. **(C)** Anthocyanin contents in *PuMYB123-like*-OE and *PuMYB123-like*-RNAi pear calli after light treatment (units: A530/g of fresh weight). **(D)** Phenotype of *Arabidopsis* seedlings ectopic expressing *PuMYB123-like* (scale bar, 2 mm). Wild-type *Col 0* was used as the negative control. *Arabidopsis* seedlings were grown on 1/2 MS medium for five days. **(E)** PCR pattern of wild-type pear calli and *PuMYB123-like* transgenic pear calli lines using transgene-specific primers. *AtPP2A* was amplified as an internal control. **(F)** Anthocyanin contents in transgenic *Arabidopsis* lines (units: A530/g of fresh weight). **(G)** The hypocotyl length of transgenic *Arabidopsis* seedlings. **(H)** Expression levels of anthocyanin-related genes in transgenic *Arabidopsis* lines. Data are presented as means ± s.d. of 3 biological replicates **(B)**, **(C)**, **(F)** and **(H)** and 15 biological replicates **(G)**. Asterisks indicate significant differences compared with wild type pear calli **(B)** and **(C)** or wild type *Arabidopsis* **(F-H)**, (two-tailed Student’s *t-test*, \**P* < 0.05, \*\**P* < 0.01), all *P* values are shown in Supplemental Data Set S7.

### Identification of PuHB40 and its response to ROS

To identify the TFs that respond to ROS and regulate *PuMYB123-like* expression under high-light stress conditions, we analyzed our RNA-seq data for the high-light stress- and DMTU-treated pear seedlings. A total of 11,608 differentially expressed transcripts (DETs) were identified by pairwise comparisons of mock, high-light stress, and high-light stress supplemented with DMTU treatments at each time point. The DETs were further categorized into 20 clusters according to their expression patterns using Mfuzz (Supplemental Fig. S7). The DETs in cluster 14 were upregulated under high-light stress conditions, but downregulated by the DMTU treatment, indicating the expression of the corresponding genes may increase in response to high-light stress-induced ROS accumulation (Fig. 4A). In addition, 60 genes that were in both the “purple” module (Supplemental Data Set S5) and cluster 14 (Supplemental Data Set S6) were identified (Fig. 4B). The transcription of these genes increased following ROS accumulation. Moreover, they were co-expressed with anthocyanin biosynthetic genes. Hence, we considered them as candidate genes involved in ROS-induced anthocyanin biosynthesis. These genes encoded five types of TFs (MYB, MYB-related, bHLH, HD-Zip, and WRKY). In terms of their transcription levels, *PubHLH3* was most highly correlated with *PuMYB123-like*, followed by *PuHB40* (HD-Zip TF) (Fig. 4C). Because PubHLH3 and PuMYB123-like interact to form an MBW complex that controls anthocyanin biosynthesis, PuHB40 was analyzed further. Notably, four HD-Zip TF-binding sites were detected in the *PuMYB123-like* promoter (Fig. 4D), suggesting that PuHB40 is the potential regulator of *PuMYB123-like* expression under high-light stress conditions.

**Figure 4.**
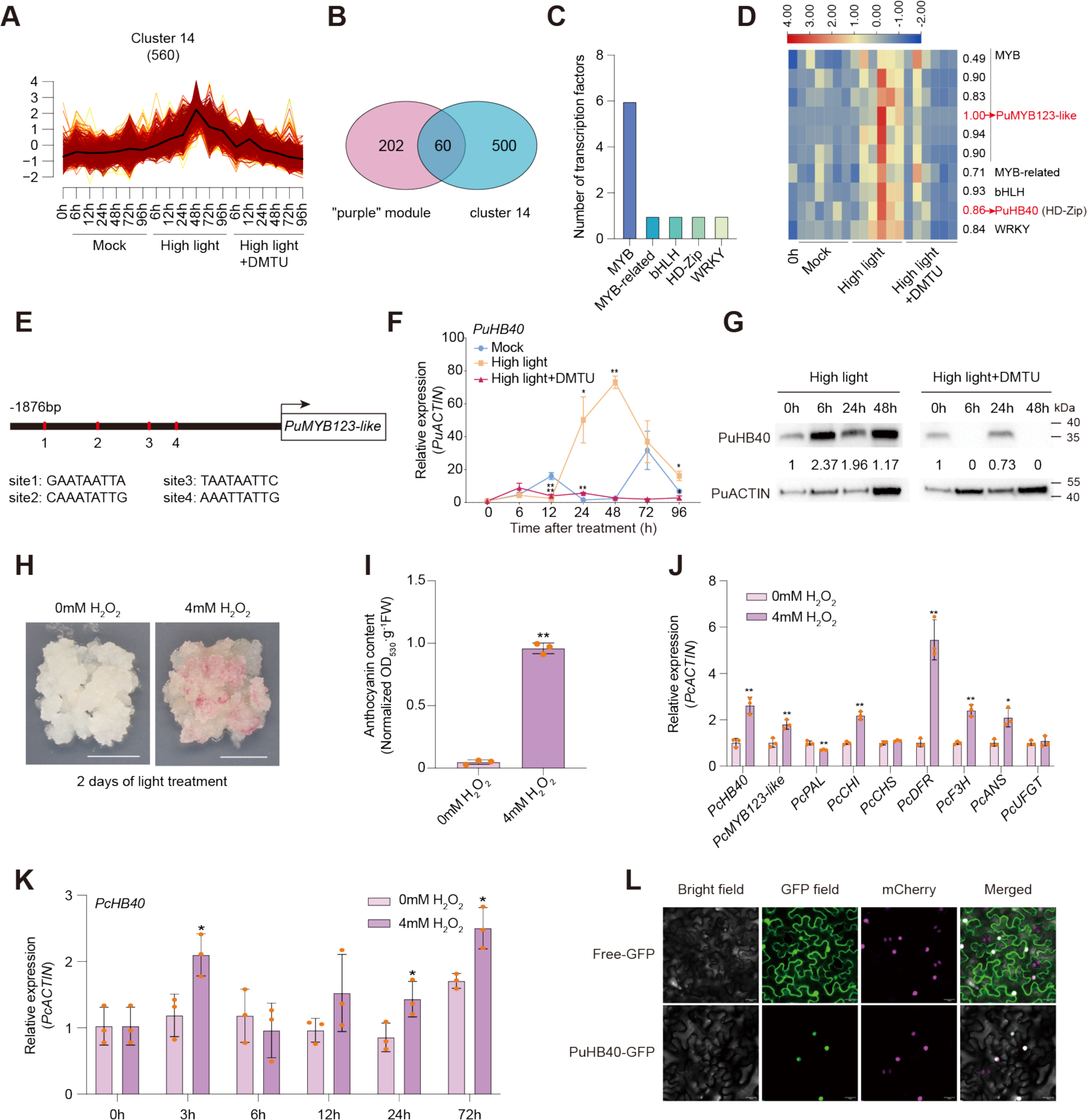
Identification of PuHB40 and its response to ROS. **(A)** The expression pattern of DETs in cluster 14 based on Mfuzz clustering. **(B)** Venn diagram of genes in “purple module” and DETs in cluster 14. **(C)** Numbers of transcription factors in overlapped genes of “purple module” and cluster 14. **(D)** Heat map presenting the expression patterns of transcription factors in overlapped genes of “purple module” and cluster 14 in response to diverse treatments. The values on the right of the heat map indicate the correlation of the expression patterns (FPKMs) between these genes and *PuMYB123-like*. **(E)** Distribution of potential binding sites of PuHB40 in *PuMYB123-like* promoter region. Vertical red lines indicate the position of site1 to site4. **(F)** The expression levels of *PuHB40* in pear seedlings under high-light treatment or high-light supplemented with DMTU treatment, and pear seedlings under normal light condition were used as Mock. *PuACTIN* was amplified as an internal control. **(G)** Changes in the PuHB40 abundance in pear seedlings under high-light treatment or high-light supplemented with DMTU treatment for 0 h, 6 h, 24 h, 48 h. The PuHB40 and PuACTIN proteins were detected by immunoblotting with an affinity-purified specific anti-PuHB40 polyclonal antibody and Anti β-Actin monoclonal antibody, respectively. **(H)** Phenotype of pear calli after 4 mM H2O2 treatment for two days (scale bar, 1 cm). Pear calli treated with 0 mM H2O2 were used as control. **(I)** Anthocyanin contents in pear calli treated with 0 mM or 4 mM H2O2 for two days (units: A530/g of fresh weight). **(J)** Expression levels of *PcHB40*, *PcMYB123-like* and anthocyanin-related genes in pear calli treated with 0 mM or 4 mM H2O2 for two days. *PcACTIN* was amplified as an internal control. **(K)** Expression levels of *PcHB40* in pear calli after 0 mM or 4 mM H2O2 treatment at 0 h, 3 h, 6 h, 12 h, 24 h, 72 h. *PcACTIN* was amplified as an internal control. **(L)** Subcellular localization of PuHB40 in *N. benthamiana* leaf cells (scale bar, 25 μm). Data are presented as means ± s.d. of 3 biological replicates. Asterisks indicate significant differences compared with Mock **(F)** or 0 mM H2O2 **(I-K)** (two-tailed Student’s *t-test*, \**P* < 0.05, \*\**P* < 0.01), all *P* values are shown in Supplemental Data Set S7.

In response to the high-light treatment, the *PuHB40* expression level gradually increased after 24 h, peaked at 48 h, and then decreased, but the DMTU treatment significantly inhibited *PuHB40* expression (Fig. 4F). Additionally, the PuHB40 content changes in response to ROS were analyzed in an immunoblot involving an affinity-purified anti-PuHB40 polyclonal antibody generated in this study (Supplemental Fig. S8). The immunoblot results showed that the PuHB40 protein content increased significantly after 6 h of the high-light treatment, after which it gradually decreased, but was maintained at a relatively high level. In contrast, the PuHB40 protein level remained low after the DMTU treatment (Fig. 4G). Hence, PuHB40 was induced by ROS after the abiotic stress treatment. To explore whether PuHB40 is directly affected by ROS, we treated pear calli with 4 mM H2O2, which resulted in a significant increase in anthocyanin biosynthesis (Fig. 4H and I). Moreover, the expression of *PcHB40* as well as the anthocyanin-structural genes *PcCHI*, *PcDFR*, *PcF3H*, and *PcANS* also increased (Fig. 4J). Following the H2O2 treatment, the *PcHB40* expression level increased after 3 h (Fig. 4K), suggestive of a direct response to H2O2.

A phylogenetic analysis showed that PuHB40 is a homolog of AT4G36740.1 (ATHB40), while a multiple sequence alignment revealed a conserved homeodomain and leucine zipper in PuHB40 (Supplemental Fig. S9, A and B). The subcellular localization analysis detected PuHB40 in the nucleus (Fig. 4L). Accordingly, PuHB40 is a transcriptional factor localized in the nucleus.

### PuHB40 promotes *PuMYB123-like* transcript level

On the basis of the presence of four possible PuHB40-binding sites in the *PuMYB123-like* promoter (binding sites 1–4), we hypothesized that PuHB40 may modulate *PuMYB123-like* expression by binding to its promoter. To test this hypothesis, we replaced the possible binding sites with TTTTTTTTT. The dual-luciferase assay detected a significant decrease in the activation of the *PuMYB123-like* promoter by PuHB40 when binding site 4 was mutated (Fig. 5A). The electrophoretic mobility shift assay (EMSA) showed that PuHB40 can directly bind to the *PuMYB123-like* promoter fragment containing binding site 4 (Fig. 5B). The chromatin immunoprecipitation (ChIP) assay revealed the specific enrichment of PuHB40 in the *PuMYB123-like* promoter region harboring binding site 4 (Fig. 5C). Thus, PuHB40 likely activates *PuMYB123-like* expression by binding to site 4 in the *PuMYB123-like* promoter.

**Figure 5.**
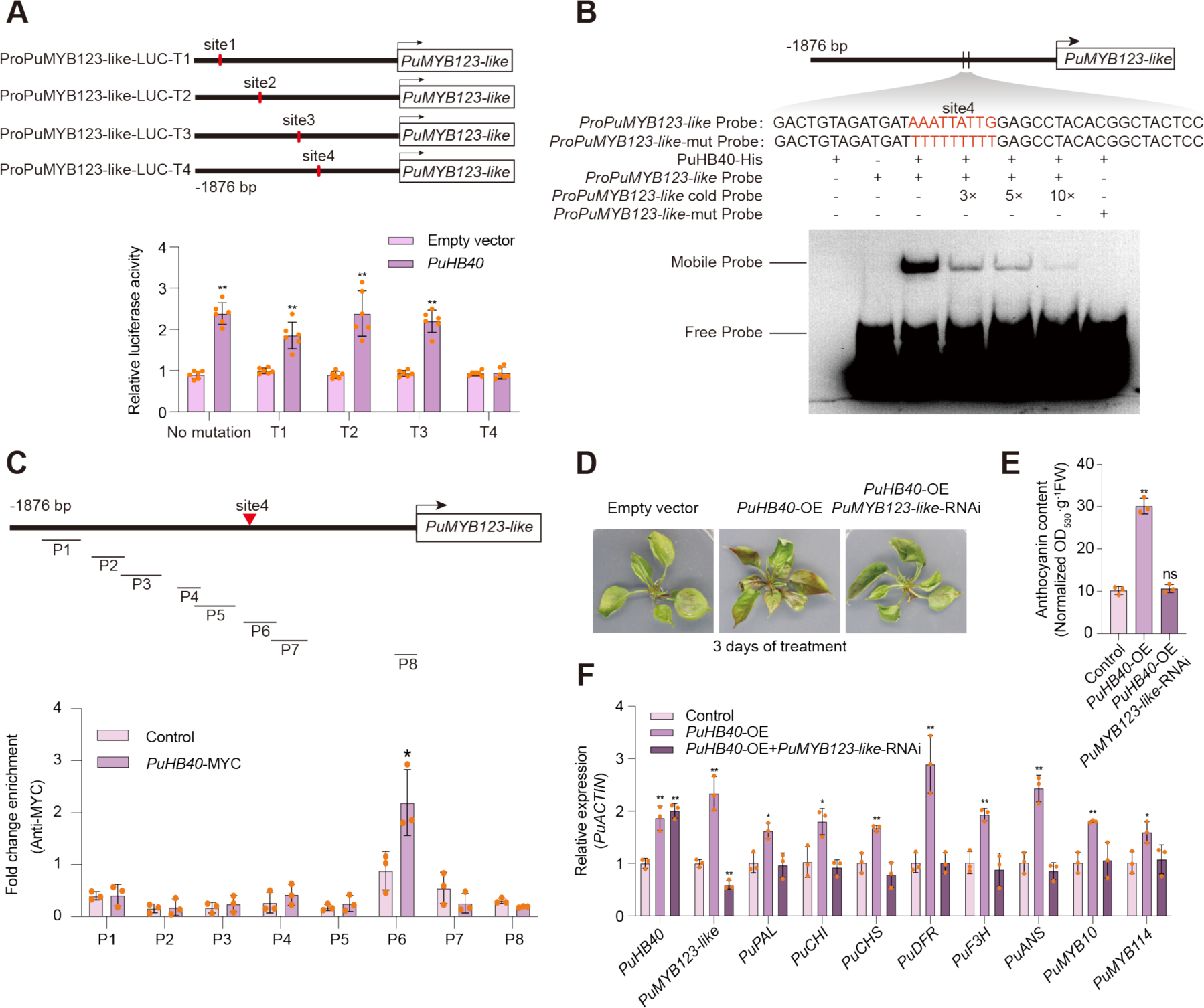
PuHB40 promotes *PuMYB123-like* transcription. **(A)** Effect of the mutation of potential PuHB40 binding sites on the activation of *PuMYB123-like*. Four potential binding sites (site1 to site4) were mutated into TTTTTTTTT, respectively. The promoters of *PuMYB123-like* with or without potential binding site mutation were cloned into the pGreenII 0800-LUC (firefly luciferase) vector, and the full-length CDS of *PuHB40* was cloned into the pGreenII 0029 62-SK vector. The empty vector of pGreenII 0029 62-SK was used as control. **(B)** The EMSA confirmed that PuHB40 bound to the site4 in the *PuMYB123-like* promoter directly. The *ProPuMYB123-like* probe was the biotin-labeled fragment of *PuMYB123-like* promoter containing site4 (AAATTATTG), whereas the cold probe was an unlabeled probe (3-, 5-, and 10-fold molar excess). *ProPuMYB123-like*-mut probe was the same as the *ProPuMYB123-like* probe but with site4 (AAATTATTG) mutated to TTTTTTTTT. **(C)** ChIP-qPCR assay results showed the binding of PuHB40 to the *PuMYB123-like* promoter *in vivo*. Cross-linked chromatin samples were extracted from *PuHB40*-MYC overexpressing pear calli with 3 biological replicates and precipitated with an anti-MYC antibody. Eluted DNA was used to amplify *PuMYB123-like* promoter fragments and eight regions (P1 to P8) were examined, among which P6 containing site4. Pear calli overexpressing the *MYC* sequence were used as a negative control. Three biological replicates of pear calli were used in the ChIP assay. **(D)** Transient overexpression of *PuHB40* with or without transient silencing of *PuMYB123-like* in pear seedlings. The full-length CDS of *PuHB40* and reverse complementary gene fragment (∼200 to 300 bp) of *PuMYB123-like* were inserted into the pCAMBIA1301 vector under the control of the 35S promoter. The plasmids were transformed into *A. tumefaciens* GV3101 and infiltrated into pear seedlings by vacuum. The infiltrated pear seedlings were firstly incubated in darkness for two days and then exposed to continuous white light for three days. Pear seedlings infiltrated with an empty pCAMBIA1301 vector were used as a control. **(E)** Anthocyanin contents in pear seedlings (units: A530/g of fresh weight). **(F)** Expression levels of *PuHB40*, *PuMYB123-like* and anthocyanin-related genes in pear seedlings. Data are presented as means ± s.d. of 3 biological replicates **(C), (E)** and **(F)** or 6 biological replicates **(A)**. Asterisks indicate significant differences compared with empty vector **(A)** or control **(C), (E)** and **(F)** (two-tailed Student’s *t-test*, \**P* < 0.05, \*\**P* < 0.01; ns, no significance, *P* > 0.05), all *P* values are shown in Supplemental Data Set S7.

To further verify the relationship between PuHB40 and *PuMYB123-like* expression, we transiently overexpressed *PuHB40* and silenced *PuMYB123-like* in pear seedlings. After a 3-day light treatment, anthocyanin accumulation was significantly greater in the *PuHB40*-OE seedlings than in the control seedlings, but the positive effect of PuHB40 on anthocyanin accumulation was significantly inhibited if *PuMYB123-like* was silenced (Fig. 5, D and E); a similar trend was detected in the expression of anthocyanin biosynthetic genes (Fig. 5F). These results confirmed that PuHB40 promotes anthocyanin biosynthesis in pear by regulating *PuMYB123-like* expression.

### PuHB40 positively regulates anthocyanin biosynthesis

To analyze the PuHB40 function related to anthocyanin biosynthesis in pear, *PuHB40*-overexpressing (*PuHB40*-OE) transgenic pear calli and *PuHB40*-RNAi pear calli were generated. After a 2-day exposure to white light, the anthocyanin contents was higher in the *PuHB40*-OE transgenic pear calli than in the control calli. Furthermore, anthocyanin accumulation was barely detectable in the *PuHB40*-RNAi pear calli (Fig. 6A-C). A qPCR analysis indicated that the anthocyanin-related genes *PcPAL*, *PcCHI*, *PcDFR*, *PcF3H*, *PcANS*, *PcUFGT*, *PcMYB10* and *PcMYB123-like* were highly expressed in the *PuHB40*-OE pear calli, whereas they were expressed at low levels in the *PuHB40*-RNAi pear calli (Fig. 6D). In addition, *PuHB40* was ectopically expressed in *A. thaliana*. Compared with the wild-type control, three independent transgenic lines ectopically overexpressing *PuHB40* had higher anthocyanin contents (Fig. 6E-G) and shorter hypocotyls (Fig. 6H). Moreover, the ectopic expression of *PuHB40* significantly increased the expression of anthocyanin-related structural genes (Fig. 6I).

**Figure 6.**
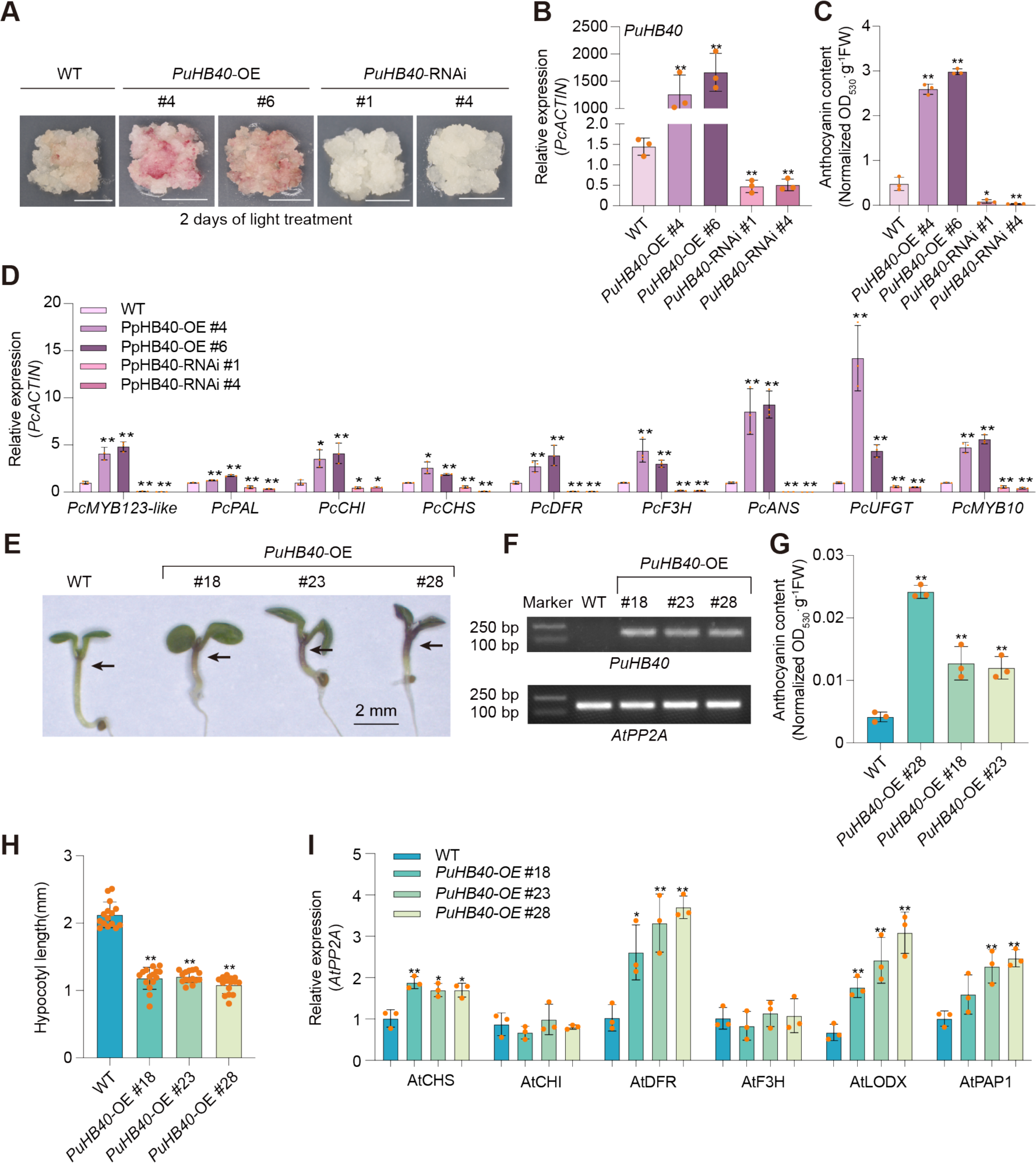
PuHB40 promotes anthocyanin biosynthesis. **(A)** Phenotype of pear calli overexpressing and silencing *PuHB40* (scale bar, 1 cm). Wild type pear calli were used as the negative control. Pear calli were incubated under continuous white light at 17°C for 3 d. **(B)** Expression levels of *PuHB40* in *PuHB40*-OE and *PuHB40*-RNAi pear calli after light treatment. *PcACTIN* was amplified as an internal control. **(C)** Anthocyanin contents in *PuHB40*-OE and *PuHB40*-RNAi pear calli after light treatment (units: A530/g of fresh weight). **(D)** Expression levels of *PcMYB123-like* and anthocyanin-related genes in *PuHB40*-OE and *PuHB40*-RNAi pear calli after light treatment. *PcACTIN* was amplified as an internal control. **(E)** Phenotype of *Arabidopsis* seedlings ectopic expressing *PuHB40* (scale bar, 2 mm). Wild-type *Arabidopsis* Col-0 was used as the negative control. *Arabidopsis* seedlings were grown on 1/2 MS medium for five days. **(F)** PCR results of wild-type pear calli and *PuHB40* transgenic pear calli lines using transgene-specific primers. *AtPP2A* was amplified as an internal control. **(G)** Anthocyanin contents in transgenic *Arabidopsis* lines (units: A530/g of fresh weight). **(H)** The hypocotyl lengths of transgenic *Arabidopsis* seedlings (units: A530/g of fresh weight). **(I)** Expression levels of anthocyanin-related genes in transgenic *Arabidopsis* lines. Data are presented as means ± s.d. of 3 biological replicates **(B-D)**, **(G)** and **(I)** or 15 biological replicates **(H)**. Asterisks indicate significant differences compared with wild type pear calli **(B-D)** or wild type *Arabidopsis* **(G-I)** (two-tailed Student’s *t-test*, \**P* < 0.05, \*\**P* < 0.01), all *P* values are shown in Supplemental Data Set S7.

To further analyze the *PuHB40* function in pear fruits, bagged mature ‘Zaosu’ and ‘Hongzaosu’ pear fruits were used for transient overexpression and VIGS experiments, respectively. The transient overexpression of *PuHB40* significantly promoted anthocyanin biosynthesis in the pear peel (Supplemental Fig. S10, A and B), while also significantly increasing the expression of anthocyanin biosynthetic genes and *PpMYB123-like* (Supplemental Fig. S10C). However, compared with the control fruit peel, the *PuHB40*-TRV fruit peel had a lower anthocyanin contents (Supplemental Fig. S10, D and E) and lower anthocyanin biosynthetic genes and *PpMYB123-like* expression levels (Supplemental Fig. S10F). Accordingly, PuHB40 may be involved in promoting anthocyanin biosynthesis. Considered together, these results imply PuHB40 functions as a positive transcriptional regulator governing anthocyanin production in pear.

### Identification of PuPP2AA2 and its interaction with PuHB40

To further explore the mechanisms through which PuHB40 regulates ROS-dependent anthocyanin biosynthesis under high-light stress, PuHB40-interacting proteins in pear seedlings were identified via a GST pull-down assay followed by a mass spectrometry analysis. Among the 36 identified proteins, we focused on the A subunit of PP2A (Supplemental Table S4). According to a phylogenetic analysis, the PP2A A subunit analyzed in the current study is most closely related to the A2 subunit of PP2A in *A. thaliana.* Thus, it was named PuPP2AA2 (Fig. 7A). During the subcellular localization analysis, PuPP2AA2 was detected in the nucleus and cytoplasm (Fig. 7B). The *PuPP2AA2* expression level was significantly lower after the high-light treatment than after the DMTU treatment (Fig. 7C). To investigate the response of PuPP2AA2 to ROS at the translational level, we transiently overexpressed the fusion construct encoding MYC-tagged PuPP2AA2 (PuPP2AA2-MYC) in pear seedlings and then exposed the plants to high-light stress conditions with or without a DMTU treatment. The PuPP2AA2 protein abundance decreased sharply under high-light conditions, but the decrease was significantly attenuated after the DMTU treatment, which presumably decreased the ROS level (Fig. 7D). Thus, PuPP2AA2 is inhibited by high-light stress-induced ROS.

**Figure 7.**
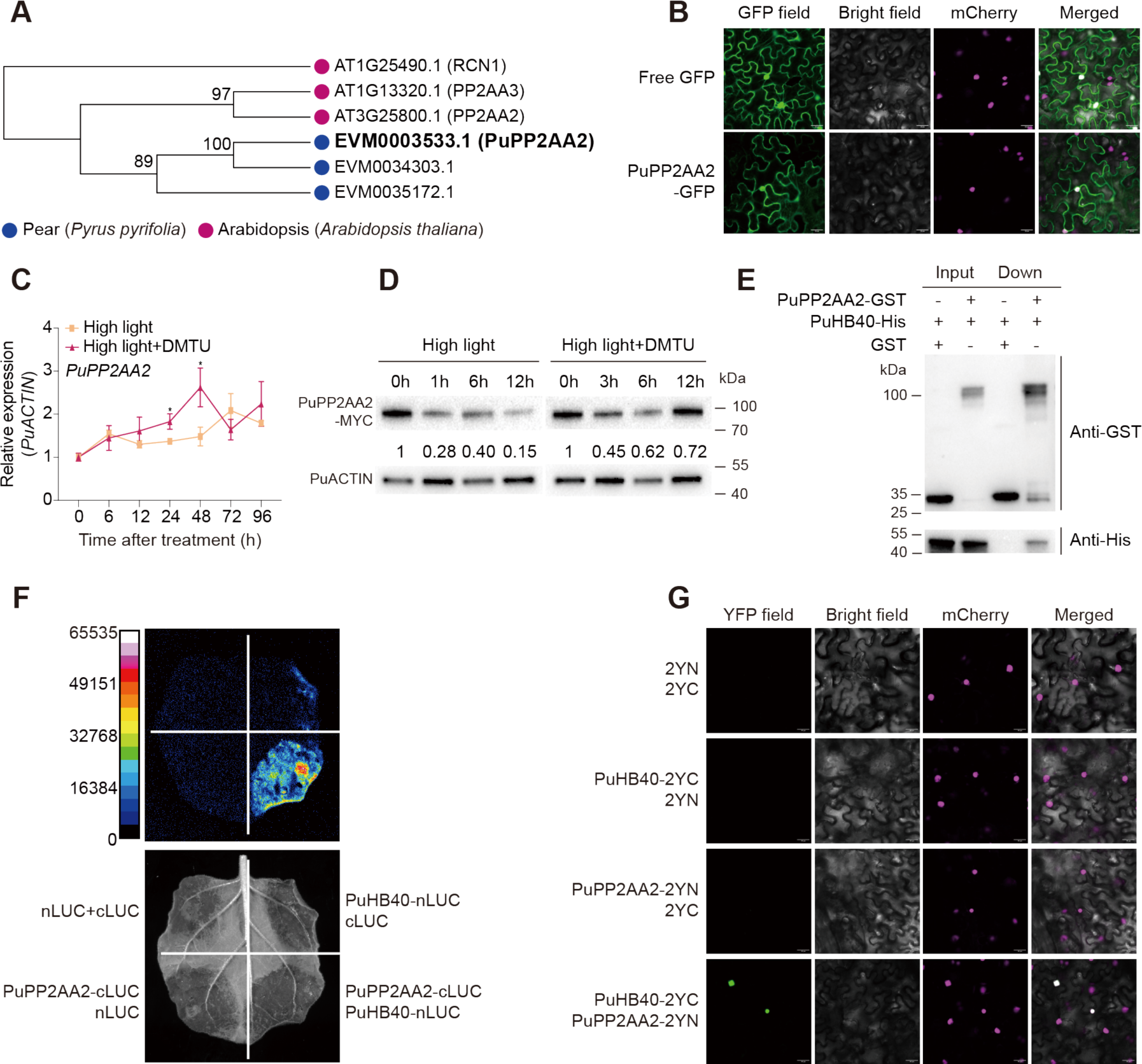
Identification of PuPP2AA2 and its interaction with PuHB40. **(A)** Phylogenetic analysis of PuPP2AA2. The phylogenetic tree was calculated using the maximum-likelihood method of MEGA X (version 10.1.8). Bootstrap values of 1000 replicates for each branch are shown. The protein sequences from pear (Pu, *Pyrus ussuriensis*, highlighted with blue dots) and Arabidopsis (*Arabidopsis thaliana*, highlighted with rose red dots). PuPP2AA2 (EVM0003533.1) is bolded and enlarged. **(B)** Subcellular localization of PuPP2AA2 in *N. benthamiana* leaf cells (scale bar, 25 μm). **(C)** The expression levels of *PuPP2AA2* in pear seedlings under high-light treatment or high-light supplemented with DMTU treatment. *PuACTIN* was amplified as an internal control. **(D)** Changes in the PuPP2AA2 abundance in pear seedlings. Pear seedlings with the transient overexpression of MYC-tagged *PuPP2AA2* were exposed to high-light treatment or high-light supplemented with DMTU treatment for 0 h, 3 h, 6 h, 12 h. The PuPP2AA2 and PuACTIN proteins were detected by immunoblotting with anti-MYC monoclonal antibody and Anti β-Actin monoclonal antibody, respectively. **(E)** The interaction between PuPP2AA2 and PuHB40 using the pull-down assay. PuHB40-His was incubated with GST or PuPP2AA2-GST. The interactive proteins were precipitated by GST-Trap agarose beads and analyzed by immunoblotting with an anti-GST or anti-His antibody. **(F)** The interaction between PuPP2AA2 and PuHB40 was detected by luciferase complementation imaging assays. The pseudocolour scale bar indicates the range of luminescence intensity. **(G)** Physical association between PuPP2AA2 and PuHB40 confirmed by a BiFC assay. Fluorescence indicates positive interactions (scale bar, 25 μm). Data are presented as means ± s.d. of 3 biological replicates. Asterisks indicate significant differences compared with high-light stress supplemented with DMTU treatment (two-tailed Student’s *t-test*, \**P* < 0.05), all *P* values are shown in Supplemental Data Set S7.

The GST pull-down assay showed that PuHB40 can interact with PuPP2AA2 *in vitro* (Fig. 7E), whereas the LCI and bimolecular fluorescence complementation (BiFC) assays reflected the interaction between PuHB40 and PuPP2AA2 *in vivo* (Fig. 7, F and G). Taken together, these data confirmed that PuHB40 interacts with the A2 subunit of PP2A, PuPP2AA2, *in vitro* and *in viv*o.

### PuPP2AA2 inhibits PuHB40-dependent PuMYB123-like trans-activation by dephosphorylating PuHB40

To clarify whether the interaction between PuPP2AA2 and PuHB40 affects the phosphorylation status of PuHB40, we transiently overexpressed the fusion construct encoding GFP-tagged PuHB40 (PuHB40-GFP) in *Nicotiana benthamiana* leaves, which were treated with cantharidin (i.e., a widely used PP2A inhibitor) (Honkanen 1993; Ren et al. 2022). We also extracted proteins from the *N. benthamiana* leaves and separated them in a Phos-tag gel and detected the fusion protein using an anti-GFP antibody. One of the two bands detected in the gel corresponded to the phosphorylated form (P+) of PuHB40-GFP, which migrated more slowly than the non-phosphorylated form (P−) of PuHB40-GFP. Moreover, the P+:P− ratio increased after the cantharidin treatment (Fig. 8A), indicating that the phosphorylation level increased after PP2A activity was inhibited. To further investigate whether PP2A can directly dephosphorylate PuHB40, we purified the MYC, PuHB40-GFP, and PP2A complex from pear seedlings that were transiently overexpressing the constructs encoding MYC, PuPP2AA2-MYC, and PuHB40-GFP using MYC-Trap and GFP-Trap agarose beads. Because PuPP2AA2 is a structural subunit that is tightly associated with the B and C subunits of the PP2A holoenzyme, other PP2A subunits may be co-purified in the PuPP2AA2-MYC immunoprecipitation complex (Zhou et al. 2004; Ren et al. 2022). Thus, the PP2A complex co-purified by PuPP2AA2-MYC may contain all of the subunits and have phosphatase activity (Ren et al. 2022). We incubated PuHB40-GFP with the MYC and PP2A complex, after which different PuHB40-GFP isoforms were detected using the anti-PuHB40 antibody for the Phos-tag analysis. The incubation with the PP2A complex led to the dephosphorylation of PuHB40 and a decrease in the P+:P− ratio from 0.57 to 0.27 (Fig. 8B), implying PP2A can directly dephosphorylate PuHB40. To confirm that the phosphatase activity of PP2A was responsible for the decrease in the phosphorylation of PuHB40, we deleted the last two tandem repeats (i.e., 14 and 15) of the HEAT motif in the C-terminal of PuPP2AA2. The mutated sequence, which was named PuPP2AA2D, was unable to bind to the B and C subunits of PP2A, thereby preventing the formation of a functional holoenzyme (Ruediger et al. 1994). The subcellular localization analysis showed that PuPP2AA2D was distributed in the nucleus as well as the cytoplasm, similar to the distribution of PuPP2AA2 (Fig. 8C). The LCI and BiFC assays demonstrated that the deletion of tandem repeats 14 and 15 did not affect the interaction between PuPP2AA2D and PuHB40 (Fig. 8, D and E).

**Figure 8.**
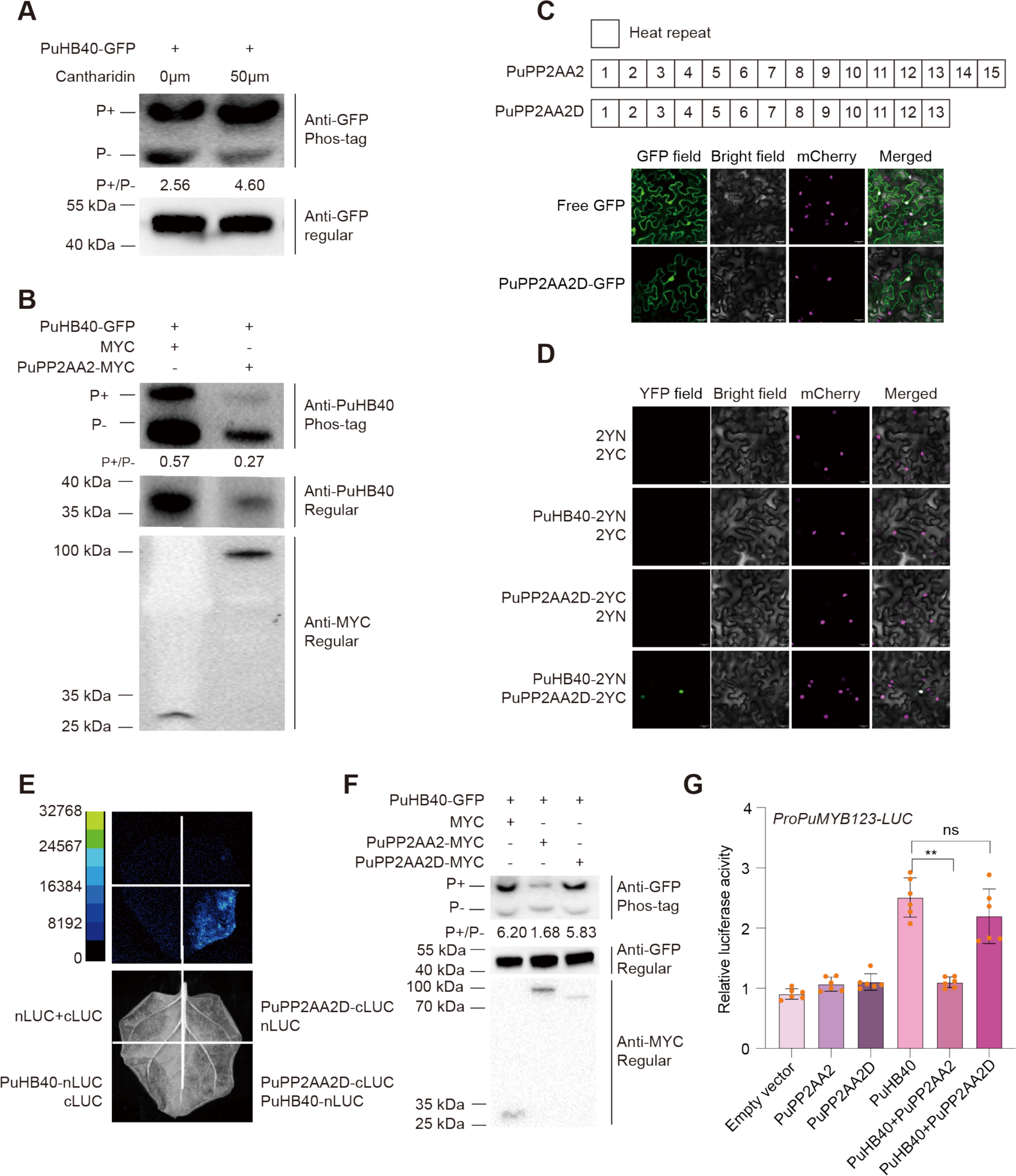
PuHB40 was dephosphorylated by PP2A phosphatase. **(A)** Phos-tag analysis of phosphorylation of PuHB40 after cantharidin treatment. *A. tumefaciens* GV3101 containing PuHB40-GFP plasmid was infiltrated into *N. benthamiana* leaves by injection and 50 μM cantharidin was injected into the same position as the agrobacteria after 24 h. *N. benthamiana* leaves injected with DMSO of corresponding concentration were used as negative control. Protein samples were extracted from *N. benthamiana* leaves after 24 h and were separated in a Phos-tag gel (upper panel), and PuHB40-GFP with different phosphorylated status was detected using the anti-GFP antibody. Total PuHB40-GFP proteins were also analyzed by SDS-PAGE without Phos-tag (indicated as “Regular”), and were detected using anti-GFP antibody (lower panel). P+ and P-indicate phosphorylated and non-phosphorylated form of PuHB40, respectively. The intensity of the bands was measured with the Image J software and the ratio of P+/P− bands was calculated. **(B)** Immunoblotting analysis showing that the PP2A complex directly dephosphorylates PuHB40 *in vitro*. MYC, PP2A complex, and PuHB40-GFP were immunopurified from pear seedlings transiently overexpressing MYC, PuPP2AA2-MYC, and PuHB40-GFP using MYC-Trap or GFP-Trap, respectively. After incubation of PP2A complexes with PuHB40-GFP (MYC with PuHB40-GFP was set as control), protein samples in the reaction system were separated in Phos-tag gel (upper panel) and detected using the anti-PuHB40 antibody. The intensity of the bands was measured with the Image J software and the ratio of P+/P− bands was calculated Total proteins in the reaction system were analyzed by SDS-PAGE without Phos-tag (indicated as “Regular”). PuHB40-GFP, MYC and PuPP2AA2-MYC were set as the loading controls and detected by the anti-PuHB40 (middle panel) or anti-MYC (lower panel) antibody, respectively. **(C)** Subcellular localization of PuPP2AA2D in *N. benthamiana* leaf cells (scale bar, 25 μm). PuPP2AA2D is PuPP2AA2 with the deletion of heat repeats 14 and 15 in the C-terminal. **(D)** The interaction between PuPP2AA2D and PuHB40 confirmed by a BiFC assay. Fluorescence indicates positive interactions (scale bar, 25 μm). **(E)** The interaction between PuPP2AA2D and PuHB40 was detected by luciferase complementation imaging assays. The pseudocolour scale bar indicates the range of luminescence intensity. **(F)** Immunoblotting analysis showing that PP2A dephosphorylates PuHB40 *in vivo*. Protein extracts from *N. benthamiana* leaves co-infiltrated PuHB40-GFP with MYC, PuPP2AA2-MYC, or PuPP2AA2D-MYC were separated in Phos-tag gel (upper panel) and detected using the anti-GFP antibody (MYC with PuHB40-GFP was set as control). The intensity of the bands was measured with the Image J software and the ratio of P+/P− bands was calculated. Total PuHB40-GFP, MYC, PuPP2AA2-MYC and PuPP2AA2D-MYC proteins were also analyzed by SDS-PAGE without Phos-tag (indicated as “Regular”). PuHB40-GFP, MYC, PuPP2AA2-MYC and PuPP2AA2D-MYC were set as the loading controls and detected by the anti-GFP (middle panel) or anti-MYC (lower panel) antibody, respectively. **(G)** Dual-luciferase assay showed that dephosphorylation of PuHB40 by PP2A significantly inhibited the activation of *PuMYB123-like* promoter by PuHB40. The *PuMYB123-like* promoter was cloned into the pGreenII 0800-LUC (firefly luciferase) vector, and the full-length CDS of *PuHB40*, *PuPP2AA2* and *PuPP2AA2D* were cloned into the pGreenII 0029 62-SK vector. The PuHB40 was used as control. Data are presented as means ± s.d. of 6 biological replicates **(G)**. Asterisks indicate significant differences compared PuHB40 **(G)** (two-tailed Student’s *t-test*, \*\**P* < 0.01; ns, no significance, *P* > 0.05), all *P* values are shown in Supplemental Data Set S7.

We transiently coexpressed the fusion constructs encoding PuHB40-GFP and PuPP2AA2-MYC as well as the fusion constructs encoding PuHB40-GFP and MYC-tagged PuPP2AA2D (PuPP2AA2D-MYC) in *N. benthamiana* leaves. Different isoforms of PuHB40-GFP were detected by the Phos-tag analysis involving the anti-GFP antibody. Furthermore, the extent of the phosphorylation of PuHB40 decreased significantly after the overexpression of *PuPP2AA2* (i.e., significant decrease in the P+:P− ratio from 6.20 to 1.68), but there was no significant change in the phosphorylation of PuHB40 after the overexpression of *PuPP2AA2D* (i.e., non-significant decrease in the P+:P− ratio from 6.20 to 5.83) (Fig. 8F). These results indicate PP2A can dephosphorylate PuHB40 *in planta*. Hence, PP2A can dephosphorylate PuHB40 *in vivo* and *in vitro*. Finally, to determine whether PuHB40 activity is influenced by its phosphorylation status, the PuHB40-induced transcription of *PuMYB123-like* was determined in a dual-luciferase assay. The presence of PuPP2AA2 abolished the ability of PuHB40 to activate *PuMYB123-like* transcription. However, the lack of PP2A phosphatase activity had no significant effect on PuHB40-activated *PuMYB123-like* transcription (Fig. 8G). Therefore, PuHB40 may interact with PuPP2AA2, with the resulting dephosphorylation by PP2A affecting whether PuHB40 can activate the transcription of *PuMYB123-like*.

### PP2A attenuates PuHB40-induced anthocyanin biosynthesis

To elucidate the contribution of PuPP2AA2 to anthocyanin biosynthesis in pear, we generated two *PuPP2AA2*-overexpressing transgenic pear calli lines (*PuPP2AA*2-OE #5 and #6), which were subsequently exposed to white light for three days. The anthocyanin contents were much lower in *PuPP2AA2*-OE #5 and #6 than in the wild-type pear calli (Fig. 9, A and B). The qPCR analysis indicated that the overexpression of *PuPP2AA2* downregulated the expression of anthocyanin-related genes (Fig. 9C). We also transiently silenced and overexpressed *PuPP2AA2* in pear fruit peels. At the injection site of the *PuPP2AA2*-OE fruit peels, there was almost a complete lack of red coloration and the anthocyanin contents was lower than that in the control fruit peels (Supplemental Fig. S11, A and B). Moreover, the expression levels of anthocyanin-related genes were significantly lower in the *PuPP2AA2*-OE pear fruit peels than in the control fruit peels (Supplemental Fig. S11C). In contrast, the transient silencing of *PuPP2AA2* significantly promoted anthocyanin accumulation (Supplemental Fig. S11, D and E) and increased the expression of anthocyanin biosynthetic genes (Supplemental Fig. S11F) suggestive of the negative regulatory effects of PuPP2AA2 on anthocyanin biosynthesis and the red coloration of pear fruits.

**Figure 9.**
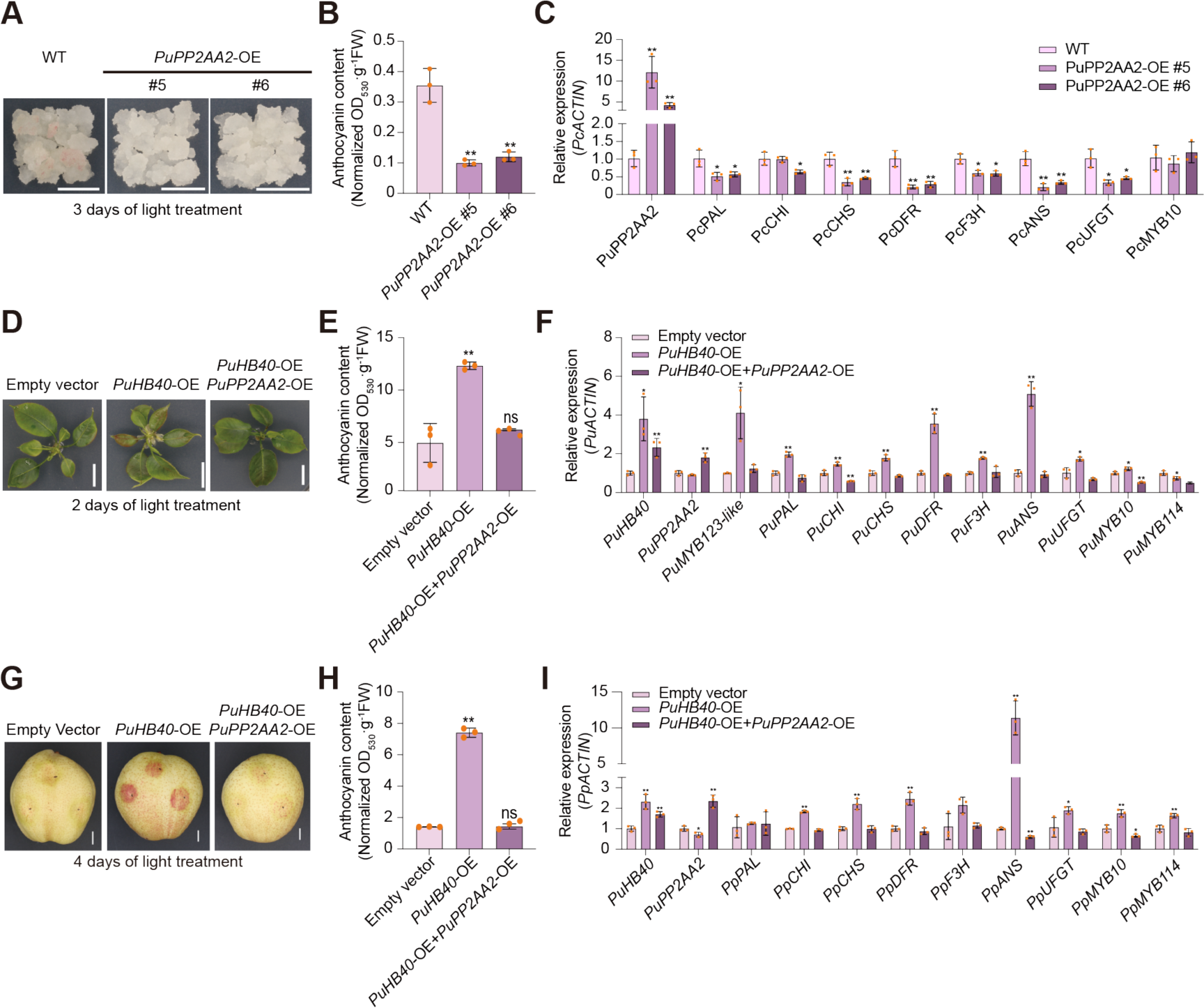
Attenuation of PuHB40-induced anthocyanin biosynthesis by PuPP2AA2. **(A)** Phenotype of pear calli overexpressing *PuPP2AA2* (scale bar, 1 cm). Wild type pear calli were used as the negative control. Pear calli were incubated under continuous white light at 17°C for 3 d. **(B)** Anthocyanin contents in *PuPP2AA2*-OE after light treatment (units: A530/g of fresh weight). **(C)** Expression levels of *PuPP2AA2* and anthocyanin-related genes in *PuPP2AA2*-OE pear calli after light treatment. *PcACTIN* was amplified as an internal control. **(D)** Transient overexpression of *PuHB40* with or without transient overexpressing of *PuPP2AA2* in pear seedlings. The full-length CDS of *PuHB40* and *PuPP2AA2* were inserted into the pCAMBIA1301 vector under the control of the 35S promoter. The plasmids were transformed into *A. tumefaciens* GV3101 and infiltrated into pear seedlings by vacuum. The infiltrated pear seedlings were firstly incubated in darkness for two days and then exposed to continuous white light for three days. Pear seedlings infiltrated with an empty pCAMBIA1301 vector were used as a control. **(E)** Anthocyanin contents in pear seedlings (units: A530/g of fresh weight). **(F)** Expression levels of *PuHB40*, *PuPP2AA2*, *PuMYB123-like* and anthocyanin-related genes in pear seedlings. **(G)** Transient overexpression of *PuHB40* with or without transient overexpressing of *PuPP2AA2* in pear fruits (scale bar, 1 cm). The full-length CDS of *PuHB40* and *PuPP2AA2* were cloned to the pGreenII 0029 62-SK vector under the control of the 35S promoter. Pear fruits were infiltrated with *A. tumefaciens* GV3101 cells containing the recombinant plasmid using a needle syringe. Fruits infiltrated with an empty pGreenII 0029 62-SK vector were used as a control. The phenotypes were examined after dark treatment for one day followed by the light treatment for four days **(H)** Total anthocyanin contents in pear fruits (units: A530/g of fresh weight). **(I)** Expression levels of *PuHB40*, *PuPP2AA2* and anthocyanin-related genes in pear fruits. Data are presented as means ± s.d. of 3 biological replicates. Asterisks indicate significant differences compared with wild type pear calli **(B)** and **(C)** or empty vector **(E), (F), (H)** and **(I)** (two-tailed Student’s *t-test*, \**P* < 0.05, \*\**P* < 0.01; ns, no significance, *P* > 0.05), all *P* values are shown in Supplemental Data Set S7.

To assess whether the phosphorylation status of PuHB40 affects its function associated with ROS-induced anthocyanin biosynthesis in pear, we transiently overexpressed *PuHB40* alone as well as with *PuPP2AA2* in pear seedlings, which were then subjected to a 2-day light treatment. Anthocyanin accumulation and the expression of anthocyanin-related genes as well as *PuMYB123-like* were greater in the pear seedlings overexpressing *PuHB40* alone than in the control seedlings. In contrast, there were no obvious differences in anthocyanin accumulation and the expression of the related genes between the control pear seedlings and the seedlings coexpressing *PuHB40* and *PuPP2AA2* (Fig. 9, D-F). Additionally, bagged mature ‘Hongzaosu’ pear fruits were used for the transient overexpression of *PuHB40* alone and with *PuPP2AA2*. Compared with the corresponding control levels, the anthocyanin contents and the expression levels of anthocyanin-related genes increased significantly in the pear fruit peels overexpressing *PuHB40* alone, but the presence of PuPP2AA2 weakened the effects of PuHB40 on anthocyanin biosynthesis and the expression of the related genes (Fig. 9, G-I). Thus, the dephosphorylation of PuHB40 by PP2A adversely affects anthocyanin biosynthesis.

## DISCUSSION

Plants growing under natural conditions over long periods are increasingly becoming more susceptible to different stresses because of the variability in environmental conditions due to global warming and climate change (Lesk et al. 2016). In plant cells, stress can induce the rapid accumulation of ROS, which are important signaling molecules that activate plant mechanisms related to defense and acclimatization (Mittler et al. 2022). High-light stress-induced ROS accumulation is closely related to photosynthetic activities. Under high-light conditions, chlorophylls are converted to their excited state following the absorption of light energy. The subsequent increase in electron transfer and energy dissipation eventually leads to the formation of highly activated triplet excited chlorophylls that readily transfer excitation energy to O2, which forms O2^−^ and H2O2 in photosystem I and ^1^O2 in photosystem II (Asada 2006; Foyer and Noctor 2009; Li et al. 2009; Mullineaux et al. 2018). Additionally, high-light stress can also induce ROS accumulation in a photosynthesis-independent manner in plant cells lacking photosynthetic pigments via a process mediated by NADPH oxidase (Xiong et al. 2021). Excessive amounts of ROS can damage plant cells and may lead to cell death (Mittler 2017) In response to a wide range of stresses and to maintain ROS homeostasis, plants have evolved a series of mechanisms, including those mediating anthocyanin accumulation.

In plants, anthocyanins are crucial antioxidants that function as ROS scavengers (Gould et al. 2002; Neill et al. 2002; Bi et al. 2014; Jiang et al. 2023). Many studies showed that anthocyanin accumulation is strongly induced by various stresses, such as high-light stress, ultraviolet radiation, and low temperatures (Steyn et al. 2002; Catalá et al. 2011; Xie et al. 2012; Nakabayashi et al. 2014; Landi et al. 2015; Xu et al. 2015). An earlier investigation involving raman spectroscopy confirmed that abiotic stress conditions increase the ROS and anthocyanin contents in the leaves of coleus plants (Altangerel et al. 2017). However, the link between ROS and anthocyanins remains unclear. Although treating *A. thaliana* seedlings with H2O2 and ROS activators promotes anthocyanin accumulation (Xu et al. 2017; Shi et al. 2018), evidence of the relationship between ROS and anthocyanin accumulation is lacking. In this study, the ROS scavenger DMTU was applied to eliminate ROS induced by various stresses, thereby serving as a reliable negative control. Stress-induced anthocyanin accumulation in pear seedlings was attenuated by DMTU (Fig. 1, A and B; Supplemental Fig. S1A), indicating that the increase in the anthocyanin contents depends on ROS, which likely function as signaling molecules linking the exposure to different stresses to changes in anthocyanin contents.

Anthocyanin biosynthesis is regulated by the MBW complex, with MYB TFs playing a major role and responding to various environmental and internal factors (Xu et al. 2015). R2R3-MYB proteins of subgroup 6 is responsible for anthocyanin biosynthesis (Dubos et al. 2010). In rosaceous plants, MYB10 TFs, which belong to subgroup 6, are essential promotors of light-induced anthocyanin biosynthesis (Takos et al. 2006; Espley et al. 2007; Feng et al. 2010). Although most of the MYB TFs promoting anthocyanin biosynthesis belong to subgroup 6, a few members of other subgroups are also involved. For example, the subgroup 2 members MdMYB9 and MdMYB11 promote jasmonic acid-induced anthocyanin biosynthesis by increasing the expression of anthocyanin-related structural genes (An et al. 2015), whereas MdMYB308L, which is a member of subgroup 4, promotes low-temperature-induced anthocyanin biosynthesis by interacting with MdbHLH33 (An et al. 2020). In the current study, we determined that PuMYB123-like, which belongs to subgroup 5, is the key MYB TF mediating anthocyanin biosynthesis via the ROS-responsive pathway. Traditionally, PuMYB10 has been considered the main MYB TF responsible for anthocyanin biosynthesis in pear (Feng et al. 2010; Bai et al. 2019a), however, its expression is similar following the high-light stress and DMTU treatments (Fig. 1M). The MYB subgroup 5 members are primarily involved in proanthocyanidin biosynthesis, but in the present study, the overexpression of *PuMYB123-like* did not affect proanthocyanidin accumulation in pear calli (Supplemental Fig. S6A). Instead, PuMYB123-like promoted anthocyanin biosynthesis by interacting with PubHLH3 (Fig. 2). Our study findings suggest that members of other MYB subgroups may also have important regulatory effects on anthocyanin biosynthesis, further clarifying the involvement of MYB TFs in ROS-associated anthocyanin biosynthesis.

According to earlier research, HD-Zip I family members are crucial regulators of plant responses to stress, especially drought and osmotic stress (Ariel et al. 2007; Harris et al. 2011). But their role in other stresses is less well known. However, their roles in responses to other stresses are less well known. In this study, one HD-Zip I family member, which was designated as PuHB40 on the basis of previously reported nomenclature (Yang et al. 2018), was identified as a key TF involved in ROS-regulated anthocyanin biosynthesis under high-light stress conditions (Fig. 4). Considering the importance of HD-Zip I family members in various stress responses, PuHB40-regulated high-light stress-induced anthocyanin biosynthesis might be a universal response to diverse stresses. The response of HD-Zip I TFs to drought stress depends on abscisic acid (ABA) (Söderman et al. 1996; Söderman et al. 1999; Olsson et al. 2004), but the relationship with ROS remains unknown. In the current study, PuHB40 responded to high-light stress in an ROS-dependent manner (Fig. 4F), implying HD-Zip I TFs can respond to other abiotic stresses (i.e., in addition to drought) through multiple signaling pathways. The stability and activity of HD-Zip proteins are regulated by phosphorylation. In apple, MdHB1 and MdHB2 are phosphorylated by SnRK2, thereby enhancing their stability and activity, which leads to increased *MdACO1* expression and ethylene biosynthesis (Jia et al. 2022). In *A. thaliana*, ATHB6 is dephosphorylated by the PP2C phosphatase ABI1, which functions as a negative regulator in the ABA signaling pathway (Himmelbach et al. 2002). The phosphorylation status of HD-Zip I TFs is mainly influenced by the ABA signaling pathway. In the present study, the phosphorylation of PuHB40 was regulated by ROS via PP2A (Fig. 8). The decrease in the extent of the phosphorylation of PuHB40 decreased the PuHB40-activated transcription of *PuMYB123-like*, ultimately decreasing ROS-induced anthocyanin biosynthesis (Fig. 9, D and G). In plants, PP2A is a ubiquitous and highly conserved serine/threonine-specific phosphatase (Farkas et al. 2007) that catalyzes dephosphorylations with diverse regulatory effects on plant stress responses (Lillo et al. 2014). In general, under salt and osmotic stress conditions, PP2A is inhibited, resulting in phosphorylated target proteins that help protect against various stresses (Bian et al. 2020; Li et al. 2020). However, the effects of ROS on PP2A phosphatase activity under abiotic stress conditions are unclear. In this study, we observed that the structural subunit of PP2A was repressed by ROS in pear (Fig. 7, C and D), suggesting that the regulatory effects of PP2A on plant stress responses may involve ROS. The A subunit of PP2A is the structural subunit that provides a scaffold that binds to the B and C subunits (Kamibayashi et al. 1992; Ruediger et al. 1994; Groves et al. 1999; Janssens and Goris 2001). In *A. thaliana*, three A subunits of PP2A have been identified (RCN1, PP2AA2, and PP2AA3). A mutation to RCN1 results in severe phenotypic changes, including stunted growth (dwarfism), abnormal embryogenesis, and sterility. This is in contrast to the mostly normal phenotypes of the *pp2aa2* and *pp2aa3* mutants (Zhou et al. 2004). Thus, it is widely believed that RCN1 is critical for the overall regulation of PP2A, which may help to explain why there have been relatively few studies on PP2AA2 and PP2AA3 (Lillo et al. 2014). In the current study, we revealed that PuPP2AA2 affects ROS-induced anthocyanin biosynthesis by dephosphorylating PuHB40 (Fig. 10), which clarifies the role of the A subunit of PP2A in plants and highlights the importance of the other A subunits.

**Figure 10.**
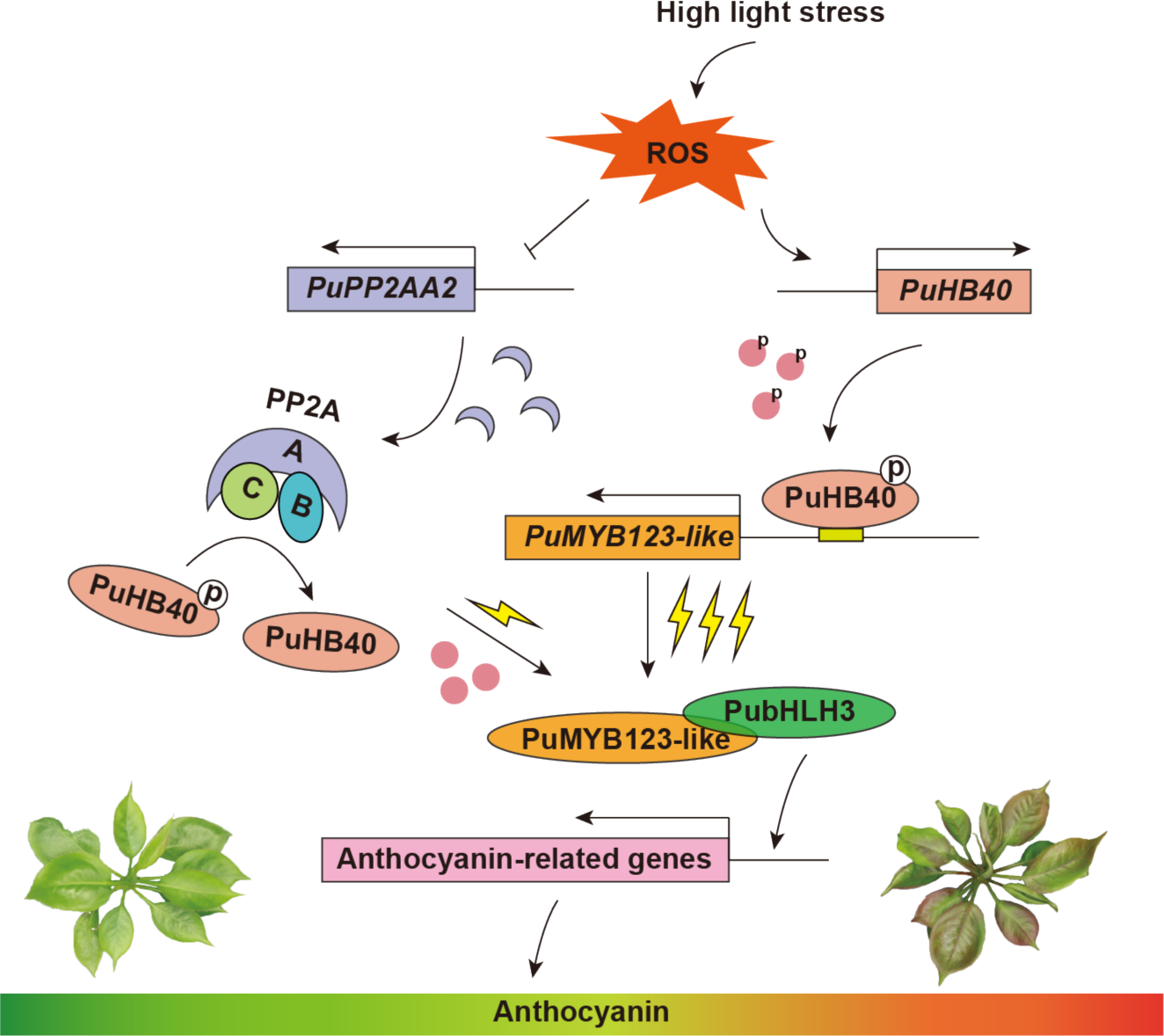
Proposed model for PuHB40-mediated ROS-dependent anthocyanin biosynthesis in pears. PuHB40 is the key transcription factor in ROS-induced anthocyanin biosynthesis. ROS promote PuHB40 expression and then positively regulate its transcriptional activity on PuMYB123-like/PubHLH3 complex to induce anthocyanin biosynthesis. The transcriptional activity of PuHB40 depends on its phosphorylation level which is regulated by PP2A. ROS maintain phosphorylation of PuHB40 at a high level, and enhance the transcriptional activity of PuHB40 on *PuMYB123-like* by suppressing the transcription of *PuPP2AA2* under high-light stress, and thus promote anthocyanin biosynthesis.

In summary, PuHB40 is the key TF regulating ROS-induced anthocyanin biosynthesis. Specifically, it is a transcriptional activator that induces PuMYB123-like/PubHLH3 complex for anthocyanin biosynthesis. The ability of PuHB40 to activate target gene transcription depends on its phosphorylation level, which is regulated by PP2A. Furthermore, ROS accumulation leads to the extensive phosphorylation of PuHB40 by suppressing the transcription of *PuPP2AA2*, thereby enhancing the PuHB40-activated transcription of *PuMYB123-like* and promoting anthocyanin biosynthesis (Fig. 10). Our study findings have elucidated the molecular mechanisms by which abiotic stresses regulate anthocyanin biosynthesis in pear, providing insights into how ROS modulate anthocyanin biosynthesis.

## MATERIALS AND METHODS

### Plant materials and treatment

The sterile seedlings of ‘Qingzhen No. 1’ pear (*Pyrus ussuriensis*) were kindly provided by Professor Ran Wang of Qingdao Agricultural University. The pear seedlings were growth at 25°C under a 16 h light/ 8 h dark photoperiod (photon flux density 50 μmol·s^-1^·m^-2^). For high-light treatment, pear seedlings were exposed to continuous LED white light (photon flux density 160 μmol·s^-1^·m^-2^) in a growth chamber. For salt and drought stress, pear seedlings were growth on solid MS medium supplemented with 30 mM sodium chloride or 2.5% polyethylene Glycol 8000 (PEG 8000) respectively and were exposed to continuous LED white light (photon flux density 50 μmol·s^-1^·m^-2^). For N, N’-dimethylthiourea (DMTU) treatment, seedlings were growth on solid MS medium supplemented with 5 mM DMTU and other conditions are same as corresponding stress treatment. After treatment, the seedlings were collected, immediately frozen in liquid nitrogen, and stored at -80°C until used.

Pear calli were generated from the flesh of ‘Clapp’s Favorite’ (*Pyrus communis*) fruitlets as previously described (Bai et al. 2019b). For H2O2 treatment, pear calli in good condition were firstly grown on the MS solid medium supplemented with 4 mM H2O2 for overnight and then were exposed to continuous white light (photon flux density 160 μmol·s^-1^·m^-2^) for two days at 17°C. The pear calli were collected after treatment and stored at -80°C until used.

### Measurement of the total anthocyanin contents

The measurement of anthocyanin contents is as previously described (Zhang et al. 2022). Briefly, 0.1 g powder of pear seedlings, pear calli or pear peels were soaked in 500 μL methanol: hydrochloric acid (99:1, v/v) solution and incubated overnight at 4°C in darkness. For pear seedlings, 250 μL chloroform and 250 μL double distilled water were added to the extracting solution for mixing and centrifuged for 10 minutes (4°C, 12000 rpm). The supernatant was measured at 530, 657 nm using the DU800 spectrophotometer (Beckman Coulter, Brea, CA, USA). The anthocyanin contents was calculated using the formula: (A530-0.25*A657)/0.1.

For pear calli and peels, the extracting solution was centrifuged at 4°C, 12000 rpm, for 10 minutes. And then, the absorbance of supernatant was measured at 530, 620, and 650 nm using the DU800 spectrophotometer (Beckman Coulter, Brea, CA, USA). The anthocyanin contents was calculated using the formula ([A530-A650]-0.2×[A650-A620])/0.1.

### Histological staining detection of ROS

H2O2 and O2^-^ were detected as previously described with minor modification (Xu et al. 2017). Briefly, the leaves of pear seedlings were dipped in DAB staining solution (for H2O2 detection) containing 1 mg·ml^-1^ DAB or NBT staining solution (for O2^-^ detection) containing 1 mg·ml^-1^ NBT. The leaves were incubated overnight in the dark with gentle shaking (25°C, 220 rpm·min^-1^). Then, the chlorophyll of leaves was removed by incubating with bleaching solution (ethanol: acetic acid: glycerol=3:1:1) at 85°C for 30 min. Finally, the leaves were kept in preservative solution (ethanol: glycerol = 4: 1) until images taken.

### RNA isolation and reverse transcription quantitative PCR

Total RNA was extracted with a modified CTAB method (Zhang et al. 2012). After genomic DNA removed, the first-strand cDNA was synthesized from 1 μg RNA with HiScript II First Strand cDNA Synthesis Kit (+gDNA wiper; Vazyme Biotech, China). qPCR was conducted using ChamQ SYBR qPCR Master Mix (Vazyme, China) on CFX Connect real-time PCR system (Bio-Rad, https://www.bio-rad.com/). The relative expression levels of genes were calculated using the 2^-△△*Ct*^ method (Livak and Schmittgen 2001) with the pear *PpActin* gene (GenBank accession no. JN684184) to normalize the expression values, and the analyses were carried out with three biological replicates. The primers for RT-qPCR are listed in Supplemental Data Set S2.

### H2O2 content quantification

The powder of pear seedlings leaves was soaked in 1 ml of 20 mM potassium phosphate buffer (pH 6.5) on ice and centrifuged (4°C, 1000 rpm), then the supernatant was transferred to new centrifuge tubes (2 ml) and additional potassium phosphate buffer was added to 1.5 ml. 100 μL plant extract was used in each reaction and the H2O2 concentration was measured with the Amplex Red hydrogen peroxide/peroxidase assay kit (Thermo Fisher Scientific, Cat. No. A22188).

### Subcellular localization analysis

Subcellular localization analysis was performed as previously described (Ni et al. 2023). the full-length complete coding sequences (CDS) (without termination codons) of the target genes (*PuMYB123-like*, *PuHB40*, *PuPP2AA2* and *PuPP2AA2D*) were inserted into pCAMBIA1300 vector to produce *Pro35S: PuMYB123-like-GFP*, *Pro35S: PuHB40-GFP*, *Pro35S: PuPP2AA2-GFP*, and *Pro35S: PuPP2AA2D-GFP*, respectively. *Agrobacterium* strain GV3101 harboring the plasmids were infiltrated into the leaves of *N. benthamiana* transgenic lines containing red fluorescent protein in the nuclei. The GFP and mCherry fluorescence signals were observed by confocal laser scanning microscopy (A1, Nikon, Tokyo, Japan). The primers used for vector construction are listed in Supplemental Data Set S1.

### Transient transformation assay in pear fruits and seedlings

‘Hongzaosu’ (also called ‘Red Zaosu’) pear fruits (*Pyrus pyrifolia* × *Pyrus communis*, genes from it are prefixed with ‘Pp’) were collected from the orchard at the Institute of Horticulture, Henan Academy of Agricultural Sciences, Henan, China. Fruits were covered with lightproof double-layered paper bags during the growing season and were harvested about 140 d after full bloom, then immediately transported to the laboratory with the bags. Transient transformation assay in pear fruits was performed as previously described (Ni et al. 2023). For the transient overexpression, the full-length CDS of *PuMYB123-like*, *PuHB40* and *PuPP2AA2* was cloned to the pGreenII 0029 62-SK vector and their expression was driven by the 35S promoter. Pear fruits were infiltrated with *A. tumefaciens* GV3101 cells containing the plasmid by a needle syringe. For VIGS assay, the gene fragment (∼200 to 300 bp) was inserted into the pTRV2 vector (gene-pTRV2) and transformed into *A. tumefaciens* GV3101. Gene-pTRV2 or empty pTRV2 (negative control) was mixed with pTRV1 in equal proportions and co-infiltrated into pear fruits by a needle syringe. After infiltration, pear fruits were exposed to continuous white light (160 μmol·m^-2^·s^-1^) and UV-B light (1.1 μmol·m^-2^·s^-1^). The fruit peel surrounding the injection sites was collected and stored at -80°C until use.

For transient transformation assay in pear seedlings, the full-length CDS of *PuHB40*, *PuPP2AA2* (for overexpression) and reverse complementary gene fragment (∼200 to 300 bp) of *PuMYB123-like* (*PuMYB123-like-RNAi*, for gene silencing) were cloned to pCAMBIA1301, and then the plasmids were transformed into *A. tumefaciens* GV3101. *A. tumefaciens* GV3101 containing *PuHB40* was mixed with empty pCAMBIA1301, *PuMYB123-like-RNAi*, or *PuPP2AA2* in equal proportions and co-infiltrated into seedlings by vacuum (0.8 kg·cm^-2^, 5 min). The seedlings were then recovered in darkness (25°C) for 1∼2 d and then transferred to a growth chamber for treatment under continuous white light (photon flux density 160 μmol·s^-1^·m^-2^). Finally, the seedlings were collected, frozen in liquid nitrogen immediately, and stored at -80°C until used. The primers used for vector construction are listed in Supplemental Data Set S1.

### Genetic transformation of pear calli

Transgenic pear calli were generated as previously described (Bai et al. 2019b). Briefly, pear calli were soaked with *A. tumefaciens* EHA105 containing different vectors (PuHB40-pCAMBIA1301, PuMYB123-like-pCAMBIA1301, PuPP2AA2-pCAMBIA1301 for overexpression and PuHB40-RNAi-pCAMBIA1301, PuMYB123-like-RNAi-pCAMBIA1301 for silencing) for 12∼15 min and then co-cultured on MS solid medium for 2 d in darkness. Positive pear calli were screened on solid MS medium supplemented with antibiotics (10 mg/L hygromycin B) under continuous darkness at 25°C for 1∼3 months and subcultured every 3 weeks. For light treatment, pear calli in good condition were exposed to continuous white light (photon flux density 160 μmol·s^-1^·m^-2^). The primers used for vector construction are listed in Supplemental Data Set S1.

### Yeast two-hybrid assays (Y2H)

The Y2H assay was carried out based with the manufacturer’s protocol of Matchmaker Gold Yeast Two-Hybrid System Kit (Takara). The primers used for vector construction are listed in Supplemental Data Set S1.

### Pull down screening coupled with LC-MS/MS

The full-length CDS of *PuHB40* was cloned to the pGEX-4T-1 vector. Then, the GST-tagged PuHB40 protein (PuHB40-GST) bound to GST-Trap agarose beads (Yeasen, Shanghai, China) were incubated with total proteins of pear seedlings at 4°C for overnight, the GST bound to GST-Trap agarose beads were used as negative control. The PuHB40-GST and GST proteins were eluted by glutathione with gravity column and subjected to immunoblotting analysis with anti-GST antibody (Cowin Biotech, China, CW0084M). The eluent solution was further analyzed by mass spectrometry.

For pull-down assay, His-tagged PuMYB123-like were incubated with GST or GST-tagged PubHLH3 in equal proportions for overnight at 4°C. Similarly, His-tagged PuHB40 were incubated with GST or GST-tagged PuPP2AA2 in equal proportions for overnight at 4°C. Then the proteins were eluted by glutathione and the eluents were subjected to immunoblotting analysis with anti-GST antibody and anti-His antibody (Cowin Biotech, China, CW0286M). The primers used for vector construction are listed in Supplemental Data Set S1.

### Luciferase complementation imaging assays

The coding sequences (without termination codons) of *PuHB40*, *PubHLH3* was inserted into pCAMBIA1300-nLUC vectors, and the CDS of *PuMYB123-like*, *PuPP2AA2* and *PuPP2AA2D* was cloned to pCAMBIA1300-cLUC vectors. The vectors were introduced into *Agrobacterium* strain GV3101 individually and *A. tumefaciens* harboring gene-nLUC were mixed with an equal volume of *Agrobacteria* containing gene-cLUC and then co-infiltrated into *N. benthamiana* leaves. Five minutes before detection, 10 mM firefly luciferin (Promega) was injected into the same position as the agrobacteria infiltration and the iluminescence was captured by ChemiDoc™XRS+ (Bio-Rad). *N. benthamiana* leaves were co-infiltrated with gene-cLUC and nLUC or cLUC and gene-nLUC as the negative control. The primers used for vector construction are listed in Supplemental Data Set S1.

### Bimolecular fluorescence complementation assay (BIFC)

BIFC assay was performed as previously described (Hu et al. 2002; Walter et al. 2004). The full-length *PuHB40* was inserted into p2YC and p2YN, while *PuPP2AA2* and *PuPP2AA2D* were cloned to p2YN and p2YN respectively. Then, *A. tumefaciens* GV3101 cells harboring gene-p2YN and gene-p2YC were mixed in equal proportions and co-infiltrated into *N. benthamiana* leaves by injection, leaves were co-infiltrated with gene-2YN and 2YC or 2YN and 2YC-gene as the negative control. The YFP signal was detected by a confocal laser scanning microscope (A1, Nikon, Tokyo, Japan) after 48 h∼72 h. The primers used for vector construction are listed in Supplemental Data Set S1.

### Dual-luciferase assay

Dual-luciferase assays were performed as previously reported (Bai et al. 2019a). The full-length CDS of *PuHB40*, *PuPP2AA2* and *PuPP2AA2D* were cloned into pGreenII 0029 62-SK vector (gene-SK) and the fragments of *PuMYB123-like* promoter were ligated into pGreenII 0800-LUC vector (promoter-LUC). All vectors were transformed into *A. tumefaciens* strain GV3101 cells harboring the pSoup vector. *A. tumefaciens* strain GV3101 containing gene-SK and promoter-LUC were injected into *N. benthamiana* leaves at a ratio of 1:10. The firefly luciferase and Renilla luciferase activities were analyzed at 48 h after the infiltration by the Dual-Luciferase Reporter Assay System (Promega) and the GloMax96 Microplate Luminometer (Promega). The primers used for vector construction are listed in Supplemental Data Set S1.

### Electrophoretic mobility shift assay

The 3’ end biotin-labeled double-stranded DNA probes were prepared by annealing 3’ end biotin-labeled oligonucleotides to the corresponding complementary strands (95 °C for 5 min and 70 °C for 20 min). The EMSA were performed using the Light-Shift^TM^ Chemiluminescent EMSA Kit (Thermo Scientific) following the manufacturer’s protocol. Briefly, 3 μg recombinant PuHB40-His protein was incubated with labeled (2 nM) or nonlabelled (6 nM, 10 nM, 20 nM) double-stranded probes at room temperature, respectively, for 20 min in binding buffer. The samples were then separated by 6.5% (w/v) native polyacrylamide gels at 4 °C and the biotin-labeled probes was captured with ChemiDoc™XRS+ (Bio-Rad). The oligonucleotide probes used in this study are listed in Supplemental Data Set S1.

### ChIP-qPCR assay

The ChIP-qPCR assays were performed as previously described (Ni et al. 2023). The pear calli transformed with the recombinant PuHB40-MYC construct and the empty construct (MYC alone) were prepared for ChIP assays. The pear calli were treated in 1% (w/v) formaldehyde under vacuum for 10 min for the cross-linking and then the chromatin was extracted via sucrose gradient centrifugation. The chromatin DNA was sonication for 30 min on ice (30 s with 30 s intervals) with the Bioruptor Plus device (Diagenode) to shear the chromatin to an average size of 200 bp to 500 bp. The sheared chromatin was incubated with Anti-MYC (Sigma-Aldrich, 05-724) at 4°C overnight and the amount of immunoprecipitated chromatin was determined by qPCR. Each ChIP assay was repeated 3 times, and the enriched DNA fragments in each ChIP sample served as 1 biological replicate for qPCR. The primers used for the ChIP-qPCR analysis are listed in Supplemental Data Set S3.

### Mn^2+^-Phos-tag SDS-PAGE

Prior to Phos-tag analysis, total protein extracted from pear leaves were pretreated by TCA precipitation and then protein samples were separated by 10% regular SDS-PAGE gel supplemented with 60 μM Phos-tag (TM) Acrylamide AAL-107 (Wako) and 120 μM MnCl2. The electrophoresis was carried out for 1.5 h using Mini Trans-Blot Electrophoretic Transfer Cell (#1703930, BIO-RAD, USA). The gel was washed by gently agitating for 10 minutes for 3 times using general transfer buffer supplemented with 10 mmol/L EDTA and then washed by general transfer buffer for 10 min. Finally, the gels were used for immunoblotting analysis.

### *In vitro* and *in vivo* protein dephosphorylation assays

For *in vitro* dephosphorylation assays, *A. tumefaciens* GV3101 harboring PuHB40-GFP, PuPP2AA2-MYC and MYC were infiltrated into pear seedlings by vacuum (0.8 kg·cm^-2^, 5 min). Then, PuHB40-GFP, PuPP2AA2-MYC and MYC were purified by immunoprecipitation from pear seedlings with GFP-Trap and MYC-Trap agarose beads. Purified PuPP2AA2-MYC was considered as PP2A complex, since other subunits of PP2A could co-immunoprecipitated with PuPP2AA2 (Zhou et al. 2004; Ren et al. 2022). Equal volume of purified PuHB40-GFP mixed with PuPP2AA2-MYC or MYC were then incubated in an oscillator (1200 g, 30 °C) for 2 h. The reaction was terminated by adding protein loading buffer and boiled at 98 °C for 10 min. Finally, the mixtures were used for immunoblotting analysis with anti-MYC monoclonal antibody (Cowin Biotech, China, CW0299M), anti-GFP polyclonal antibody (HuaBio, China, R1312-2) and anti-PuHB40 (ABclonal Technology, Woburn, USA).

For *in vivo* dephosphorylation assays, *A. tumefaciens* GV3101 harboring PuHB40-GFP was mixed with *A. tumefaciens* GV3101 harboring MYC, PuPP2AA2-MYC, PuPP2AA2D-MYC in equal proportions and co-infiltrated into *N. benthamiana* leaves by injection. The total proteins were extracted from *N. benthamiana* leaves after 48 h-72 h and subjected to TCA precipitation, immediately followed by Phos-tag analysis. For cantharidin treatment, *A. tumefaciens* GV3101 with PuHB40-GFP was infiltrated into *N. benthamiana* leaves for 24 h. Then, 50 μM cantharidin was injected into the same position as the agrobacteria infiltration and waited for 24 h. Finally, the total proteins were extracted and then used for Phos-tag analysis.

### The preparation of PuHB40-specific antibody

An affinity-purified anti-PuHB40 polyclonal antibody was generated by ABclonal Technology (Woburn, USA) with the specific fragment of *in vitro* expressed PuHB40 (60-215 aa) amplified from cDNA of pear seedling (*Pyrus ussuriensis*).

### Sequence analysis and phylogenetic analysis

Protein sequence alignments were performed using MEGA X software (Kumar et al. 2018) with default parameters. The phylogenetic tree was constructed via the maximum-likelihood method in MEGA X software. The protein sequence alignments used to construct the phylogenetic tree are provided in Supplemental File S1/S3/S5/S7, and the tree files are provided in Supplemental File S2/S4/S6/S8.

### RNA-seq data analysis

The library construction and sequencing were performed by Novogene (Beijing, China) on an Illumina Novaseq platform with a 150-bp pair-end strategy. Clean data was obtained from the raw data by eliminating low-quality reads as well as reads containing an adapter or poly-N sequences and were mapped to reference genome of ‘Cuiguan’ pear (Gao et al. 2021). The expression levels of genes were calculated based on FPKM. The DETs were identified by differential gene expression analysis tool of TBtools-II (Chen et al. 2023). The obtained *P*-values were adjusted based on Benjamini and Hochberg method, and the genes with adjusted *P*-values less than 0.05 were finally identified as differentially expressed genes (fold-change ≥ 2, false discovery rate < 0.01).

### Statistical analysis

All data in this study were analyzed using two-sided Student’s *t*-test or one-way analysis of variance (ANOVA) to determine statistical significance with GraphPad Prism v.9.0. Detailed statistical analysis data are shown in Supplemental Data Set S7.

## ACCESSION NUMBERS

The RNA sequencing data have been deposited in the NCBI Sequence Read Archive (http://www.ncbi.nlm.nih.gov/sra/), with the BioProject ID: PRJNA1098349. Sequence data of genes from this article can be found in an online pear genome database (https://ngdc.cncb.ac.cn/gwh/Assembly/18534/show): PuHB40 (EVM0020592.1), PuMYB123-like (EVM0034726.1), PuPP2AA2 (EVM0003533.1), PubHLH3 (EVM0031731.1), PubHLH33 (EVM0038434.1), PubHLH64 (EVM0003136.1), PuELIP1 (EVM0019834.1), PuELIP2 (EVM0035306.1), PuCHS (EVM0036585.1), PuCHI (EVM0024725.1), PuF3H (EVM0021098.1), PuDFR (EVM0012887.1), PuANS (EVM0001840.1), PuUFGT (EVM0026501.1), PuMYB10 (EVM0039871.1), PuMYB114 (EVM0020507.1).

## SUPPLEMENTAL DATA

**Supplemental Figure S1**. Stresses induced anthocyanin biosynthesis is ROS-dependent (Supports Figure 1).

**Supplemental Figure S2.** Identification of transcripts by the weighted gene co-expression network analysis (WGCNA) of high-light stress-treated or high-light stress supplemented with DMTU treatment in pear seedlings (Supports Figure 2).

**Supplemental Figure S3**. Expression levels of eight R2R3-MYB TFs in “purple” module in pear seedlings treated with high-light stress or high-light stress supplemented with DMTU (Supports Figure 2).

**Supplemental Figure S4.** Identification of PuMYB123-like (Supports Figure 2).

**Supplemental Figure S5.** The interaction between PuMYB123-like and PubHLHs (Supports Figure 2).

**Supplemental Figure S6.** Functional analysis of PuMYB123-like in ‘Hongzaosu’ and ‘Zaosu’ pear fruit (Supports Figure 3).

**Supplemental Figure S7.** Results of the Mfuzz clustering of 11,608 differentially expressed transcripts based on their expression patterns (Supports Figure 4).

**Supplemental Figure S8.** Detection of anti-PuHB40 polyclonal antibody specificity (Supports Figure 4).

**Supplemental Figure S9.** Identification of PuHB40 (Supports Figure 4).

**Supplemental Figure S10**. Functional analysis of PuHB40 in ‘Hongzaosu’ and ‘Zaosu’ pear fruit (Supports Figure 6).

**Supplemental Figure S11.** Functional analysis of PuPP2AA2 in ‘Hongzaosu’ pear fruit (Supports Figure 9).

**Supplemental Data Set S1.** Primers used for constructing vectors or EMSA in this study.

**Supplemental Data Set S2.** Primers for qPCR analysis in this study.

**Supplemental Data Set S3.** Primers for ChIP-qPCR analysis in this study.

**Supplemental Data Set S4.** PuHB40 possible interacting proteins were identified with GST pull-down combined with mass spectrometry.

**Supplemental Data Set S5.** The FPKM value and annotations of genes in “purple” module.

**Supplemental Data Set S6.** The FPKM value and annotations of DETs in cluster 14.

**Supplemental Data Set S7.** Detailed statistical analysis in this study.

**Supplemental File S1.** Protein sequences used to generate phylogenetic tree in Fig. 2B.

**Supplemental File S2.** Phylogenetic tree of Fig. 2B in Newick format.

**Supplemental File S3.** Protein sequences used to generate phylogenetic tree in Fig. 7A.

**Supplemental File S4.** Phylogenetic tree of Fig. 7A in Newick format.

**Supplemental File S5.** Protein sequences used to generate phylogenetic tree in Fig. S4A.

**Supplemental File S6.** Phylogenetic tree of Fig. S4A in Newick format.

**Supplemental File S7.** Protein sequences used to generate phylogenetic tree in Fig. S9A.

**Supplemental File S8.** Phylogenetic tree of Fig. S9A in Newick format.

## ACKNOWLEDGMENTS

We thank R. Wang at Qingdao Agricultural University for providing pear seedlings of ‘Qingzhen No. 1’ (*Pyrus ussuriensis*). We also thank Kunfeng Li for technical help and providing plant materials.

## FUNDING INFORMATION

This work was supported by the Zhejiang Provincial Natural Science Foundation of China under Grant No. LR22C150001, The National Natural Science Foundation of China (32072545 to YT, 32272639 to SB, 32260745 to JS), and the China Agriculture Research System of MOF and MARA.

## AUTHOR CONTRIBUTIONS

Y.T and S.B. planned the work. S.B. and L.Z. conceived and designed the experiments. L.Z. performed most experiments. L.W., S.Y. helped conduct some of the experiments. Y.F., Y.G. and J.S. helped to analyze the data. L.Z. and S.B wrote the manuscript and, Y.T., J.N. helped to revise the manuscript. All authors read and approved the final paper.

## DECLARATION OF INTERESTS

The authors declare that they have no competing interests.

## Notes

### Competing Interest Statement

The authors have declared no competing interest.

### Summary of Updates

1. We have added RNA-seq analysis data in the revised manuscript, especially the procedure for identification of PuMYB123-like and PuHB40 from the RNA-Seq data 2. Figure 1 revised.The results of NBT and DAB staining quantitative analysis have been added in Fig. 1, D and F. 3. To ensure that the work is presented in the best manner, we have revised the entire manuscript using a professional editing service. 4. Provide evidence for the reactivity and specificity of DMTU in plants. 5. Figure 10 revised. Specifically, we clearly demonstrated that ROS inhibited the transcription of PuPP2AA2 in the diagram.

