## Supplemental Figures for "Phosphorylated transcription factor PuHB40 is involved in ROS-dependent anthocyanin biosynthesis in pear exposed to high-light stress"

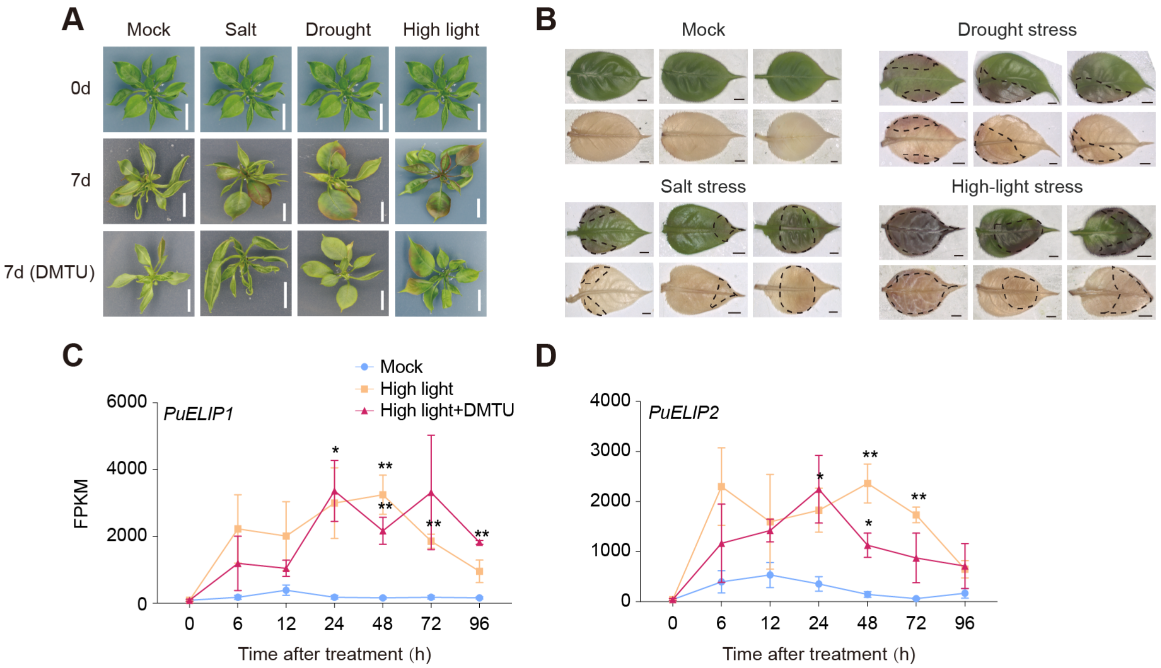


**Supplemental Figure S1. Stresses induced anthocyanin biosynthesis is ROS-dependent (Supports Figure 1). (A)** Phenotype of pear seedlings under various stress treatment (salt stress, drought stress and high-light stress) or stresses supplemented with DMTU treatment for seven days. Pear seedlings under normal growth condition were used as Mock (scale bar, 1 cm). **(B)** The detection of anthocyanin and $H_2O_2$ accumulation region in pear seedlings leaves under salt stress, drought stress and high-light stress treatment. The leaves of pear seedlings under normal growth condition were used as Mock (scale bar, 2 mm). DAB staining for $H_2O_2$ in pear seedlings leaves. The area circled in dotted lines indicates anthocyanin and $H_2O_2$ accumulation region. **(C)** and **(D)** The FPKM value of high-light stress responsive genes *PuELIP1* and *PuELIP2* in pear seedlings under high-light reatment or high-light supplemented with DMTU treatment, and pear seedlings under normal light condition were used as Mock. Data are presented as means ± s.d. of 3 biological replicates. Asterisks indicate significant differences compared with Mock group (two-tailed Student's *t-test*, $*P < 0.05$, $**P < 0.01$), all *P* values are shown in Supplemental Data Set S7.

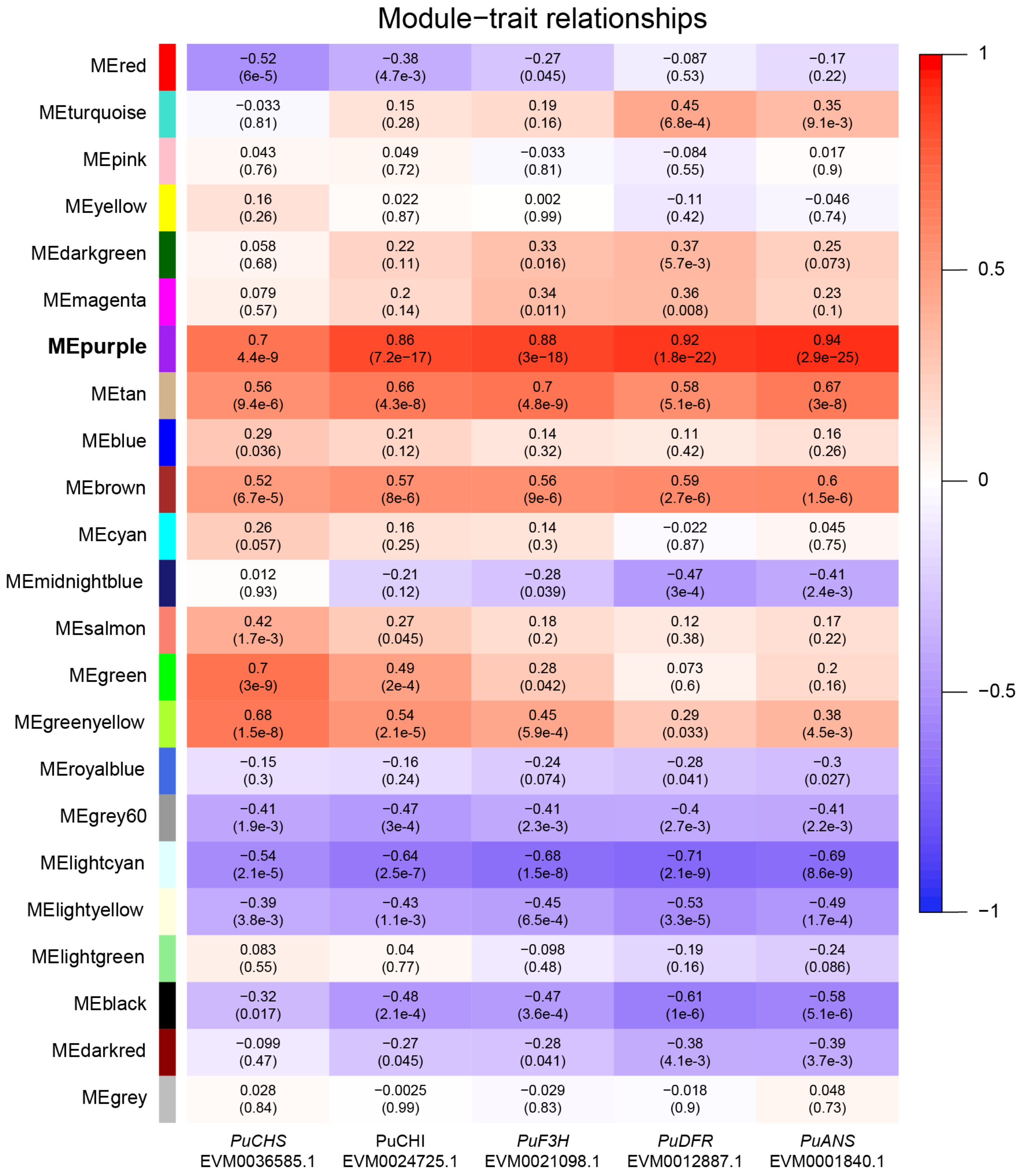


**Supplemental Figure S2. Identification of transcripts by the weighted gene co-expression network analysis (WGCNA) of high-light-treated or high-light supplemented with DMTU treatment in pear seedlings (Supports Figure 2).** Module–trait correlations and the corresponding *P*-values are in parentheses. The expression level (FPKM) of anthocyanin-related structural genes function as trait data. The left panel presents the 23 modules. The color scale on the right presents the module–trait correlations from -1 (purple) to 1 (red).

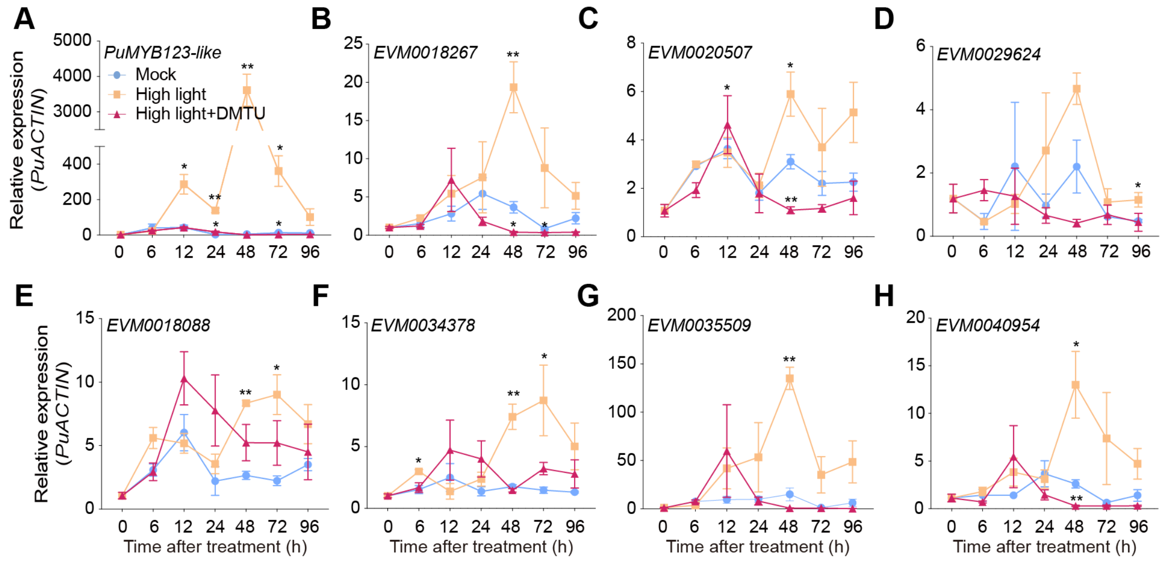


**Supplemental Figure S3. (A-H) Expression levels of eight R2R3-MYB TFs in "purple" module in pear seedlings treated with high-light or high-light supplemented with DMTU treatment (Supports Figure 2).** Pear seedlings under normal light condition were used as Mock. *PuACTIN* was amplified as an internal control. Data are presented as means ± s.d. of 3 biological replicates. Asterisks indicate significant differences compared with Mock group (two-tailed Student's *t-test*, *$P < 0.05$, **$P < 0.01$), all $P$ values are shown in Supplemental Data Set S7.

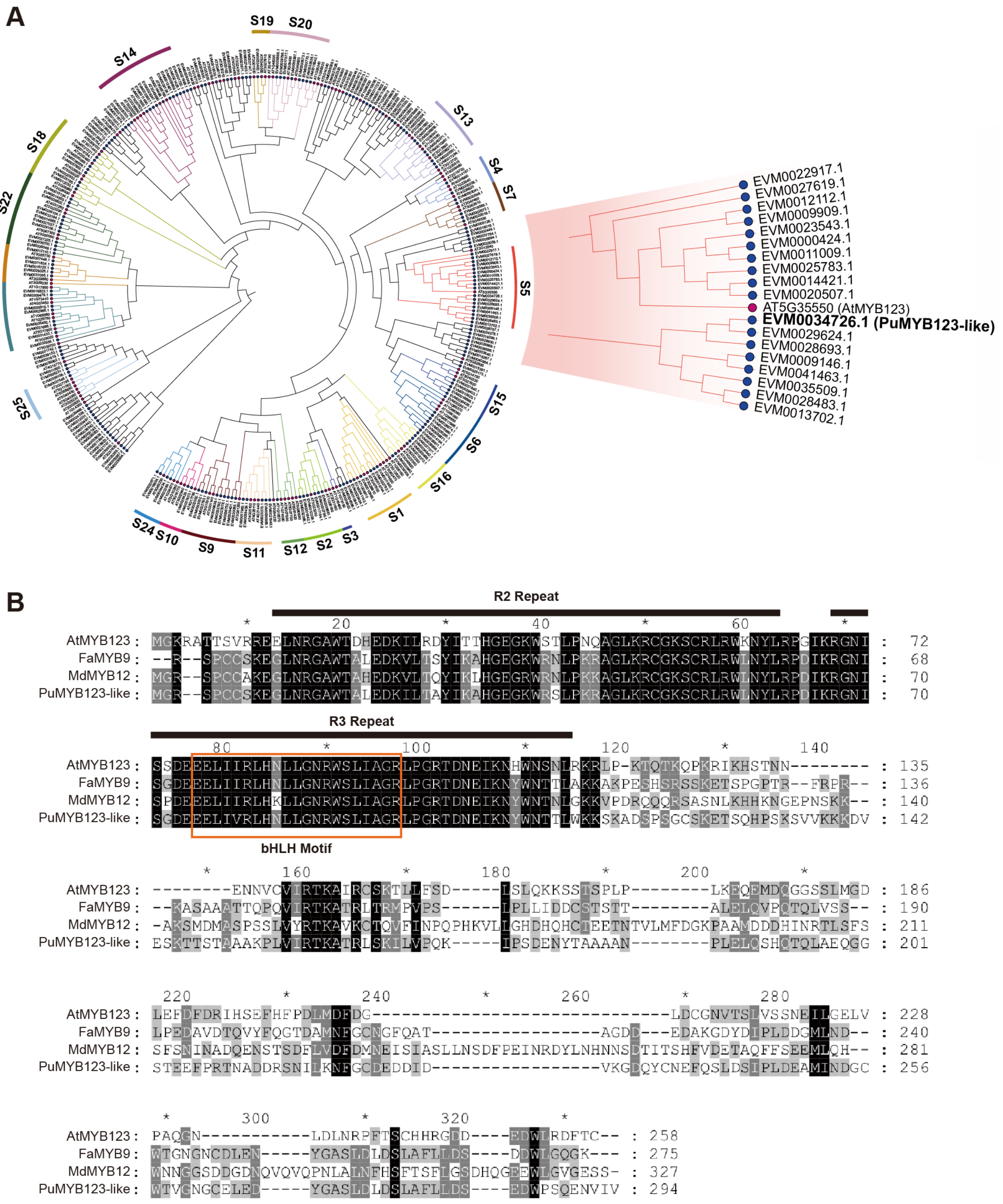


**Supplemental Figure S4. Identification of PuMYB123-like (Supports Figure 2). (A)** Phylogenetic analysis of candidate R2R3-MYB TFs from pear and *Arabidopsis*. The phylogenetic tree was generated using the maximum-likelihood method of MEGA X (version 10.1.8). Bootstrap values of 1000 replicates for each branch are shown. Different subgroups are marked with different colors respectively and S5 subgroup is enlarged on the right. The protein sequences from pear (Pu, *Pyrus ussuriensis*, highlighted with blue dots), Arabidopsis (At, *Arabidopsis thaliana*, highlighted with rose red dots). PuMYB123-like (EVM0034726.1) is bolded and enlarged. **(B)** Amino acid sequence alignment of PuMYB123-like with the homologous R2R3-MYB genes in other plants. The R2R3 DNA-binding domain is shown and the predicted motif interacting with bHLH proteins are indicated. Related proteins including *Arabidopsis thaliana* AtMYB123 (AT5G35550.1), *Fragaria × ananassa* FaMYB9 (AFL02460.1), *Malus × domestica* MdMYB12 (MDP0000887107), *Pyrus ussuriensis* PuMYB123-like (EVM0034726.1).

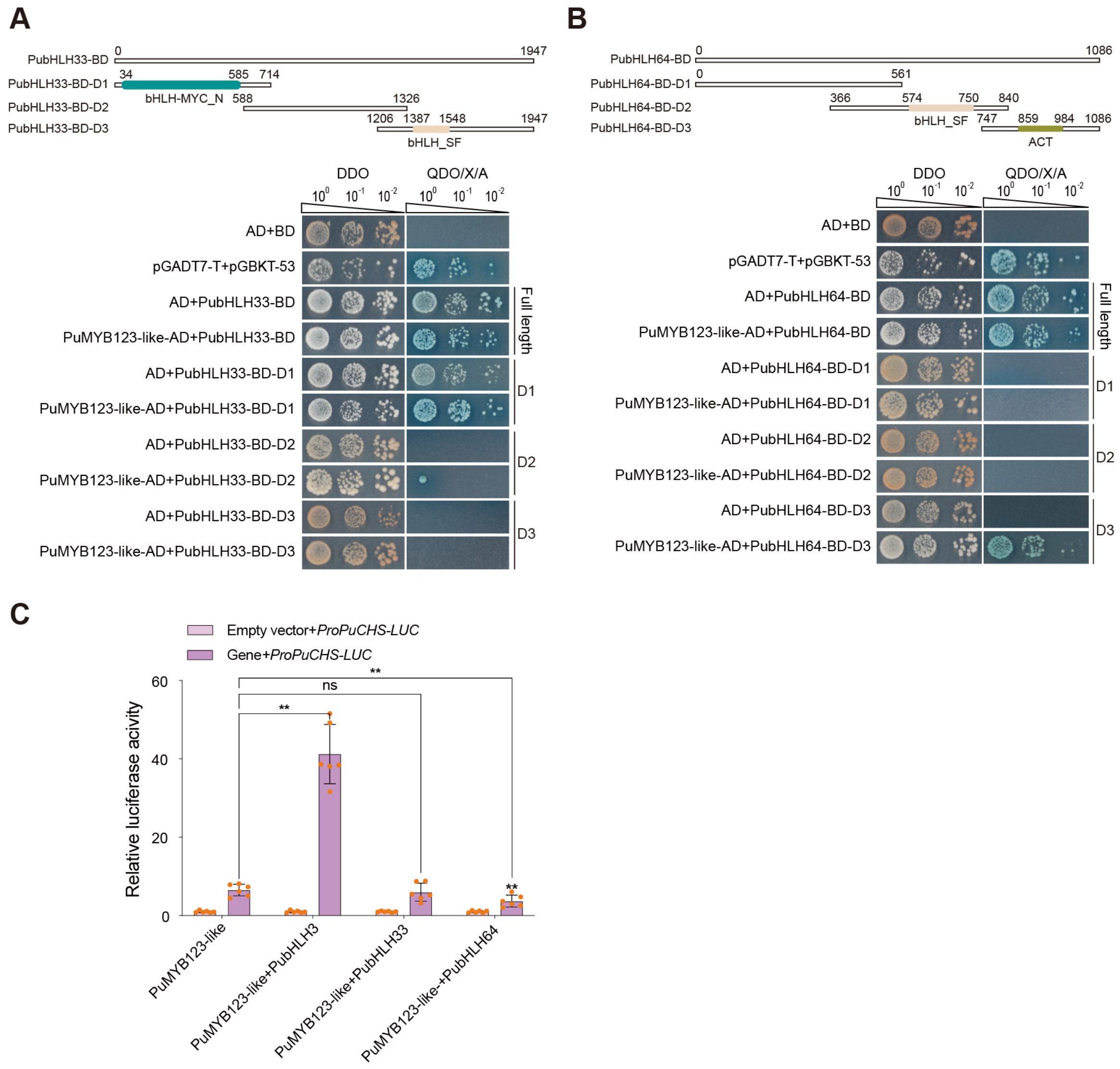


**Supplemental Figure S5. The interaction between PuMYB123-like and PubHLHs (Supports Figure 2). (A)** and **(B)** The interaction between PuMYB123-like and PubHLH33 or PubHLH64 in a yeast two-hybrid assay, with pGADT7-T and pGBKT7-53 as positive controls. As PubHLH33 and PubHLH64 showed self-activation, these two genes were divided to three fragments (D1 to D3) to test the interactions. DDO, SD/-Trp/-Leu medium; QDO/X/A, SD/-Trp/-Leu/-Ade/-His medium with X-α-gal and Aureobasidin. **(C)** Dual-luciferase assay showed that the effect of interaction between PuMYB123-like and PubHLH3, PubHLH33 or PubHLH64 on the activation of PuMYB123-like to *PuCHS* promoters. Data are presented as means ± s.d. of 6 biological replicates **(C)**. Asterisks indicate significant differences compared with PuMYB123-like **(C)** (two-tailed Student's *t-test*, $^{**}P < 0.01$; ns, no significance, $P > 0.05$), all *P* values are shown in Supplemental Data Set S7.

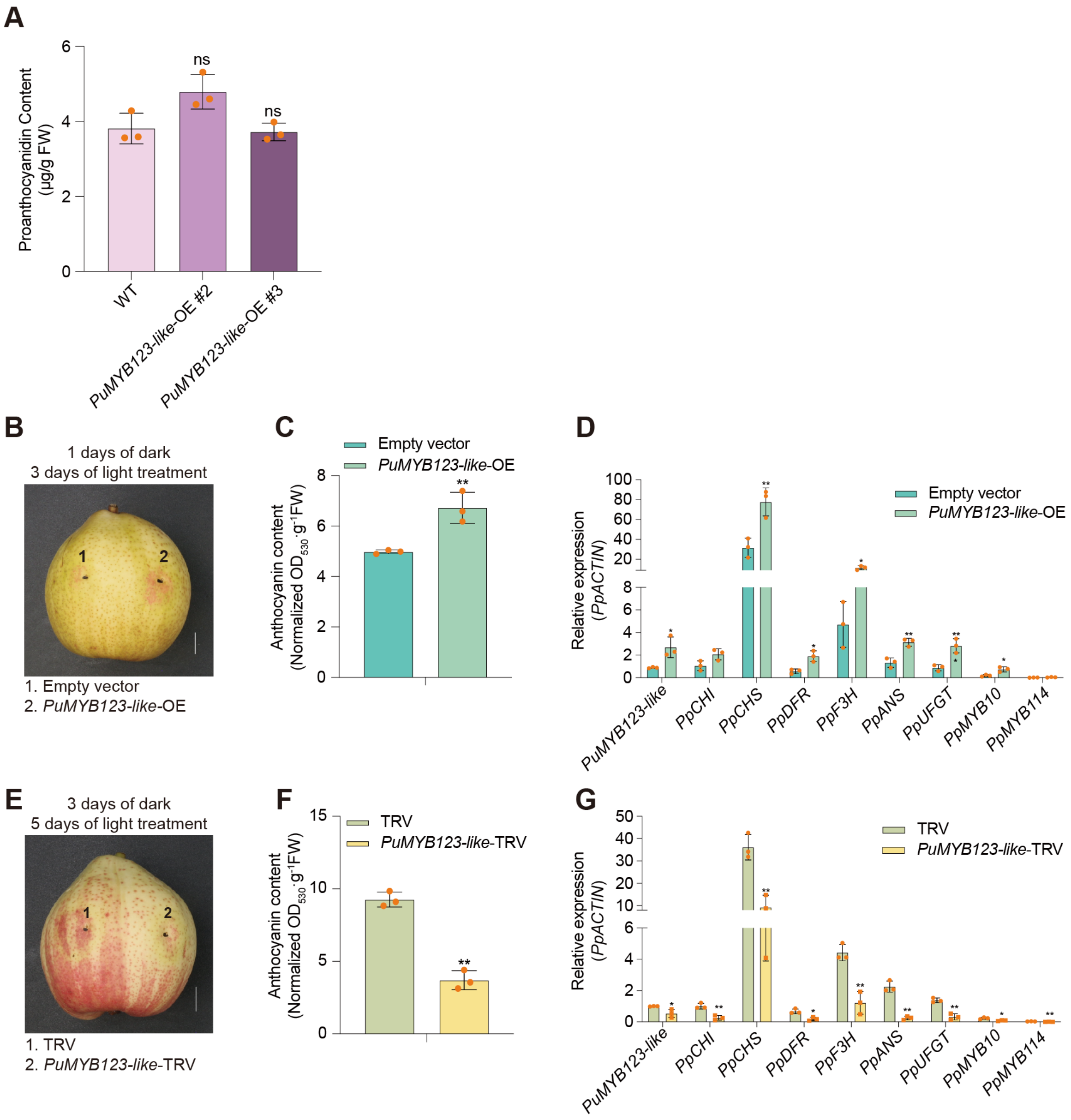


**Supplemental Figure S6. Functional analysis of PuMYB123-like in ‘Hongzaosu’ and ‘Zaosu’ pear fruits (Supports Figure 3). (A)** Proanthocyanidin contents in *PuMYB123-like*-OE pear calli. **(B)** Transient overexpression of *PuMYB123-like* in in pear fruits (scale bar, 1 cm). The full-length CDS of *PuMYB123-like* was inserted into the pGreenII 0029 62-SK vector under the control of the 35S promoter. Pear fruits were infiltrated with *A. tumefaciens* GV3101 cells containing the recombinant plasmid using a needle syringe. Fruits infiltrated with an empty pGreenII 0029 62-SK vector were used as a control. The phenotypes were examined after dark treatment for one day followed by the light treatment for three days **(C)** Total anthocyanin contents in fruits transiently overexpressing *PuMYB123-like* (units: $A_{530}$/g of fresh weight). **(D)** Expression patterns of genes related to anthocyanin biosynthesis in fruits transiently overexpressing *PuMYB123-like*. **(E)** Transient silencing of *PuMYB123-like* in fruits (scale bar, 1 cm). The empty vector (pTRV1 + pTRV2) was used as the negative control. Pear fruits were placed in darkness for three days and then treated with strong light for five days. **(F)** Total anthocyanin contents in fruits transiently silencing *PuMYB123-like* (units: $A_{530}$/g of fresh weight). **(G)**, Expression patterns of genes related to anthocyanin biosynthesis in fruits in which *PuMYB123-like* was transiently silenced. Data are presented as means ± s.d. of 3 biological replicates. Asterisks indicate significant differences compared with wild type calli **(A)**, empty vector **(C)** and **(D)** or TRV **(F)** and **(G)** (two-tailed Student's *t-test*, $*P < 0.05$, $**P < 0.01$; ns, no significance, $P > 0.05$), all $P$ values are shown in Supplemental Data Set S7.

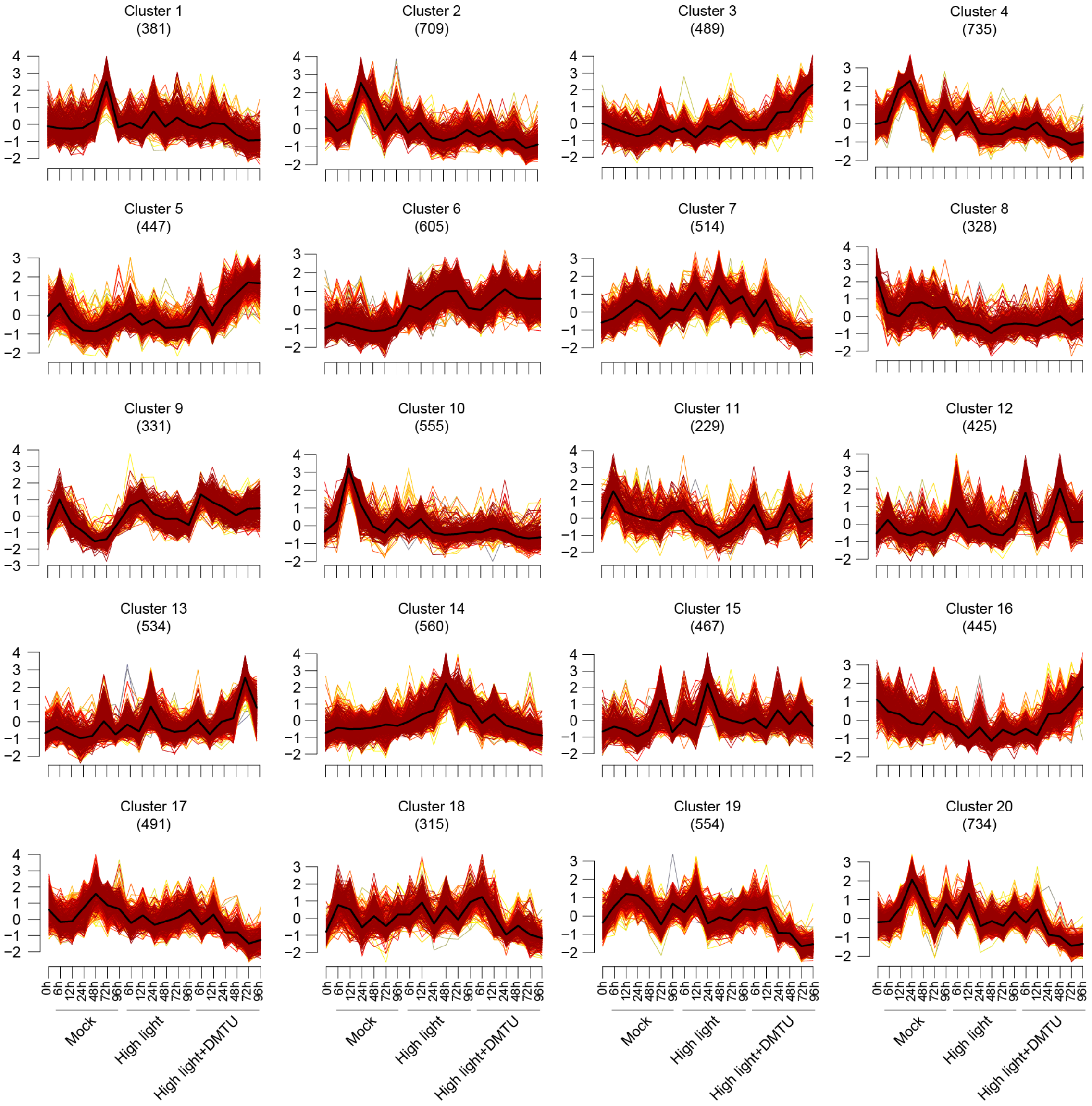


**Supplemental Figure S7. Results of the Mfuzz clustering of 11,608 differentially expressed transcripts based on their expression patterns (Supports Figure 4).**

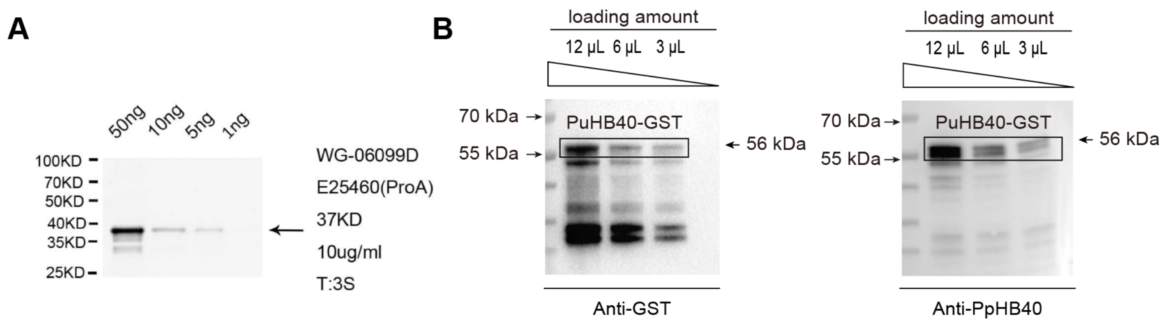


**Supplemental Figure S8. Detection of anti-PuHB40 polyclonal antibody specificity (Supports Figure 4). (A)** The arrow indicates the size of antigens pET-28a-SUMO-PuHB40 (60-215aa) targeted by anti-PuHB40. 50 ng, 10 ng, 5 ng and 1 ng antigens were used for detection, respectively. **(B)** Purified PuHB40-GST protein was used to immunoblot analysis via anti-GST and anti-PuHB40, respectively. Black arrows on the right indicate a molecular size of approximately 56 kDa corresponding to the putative PuHB40-GST.

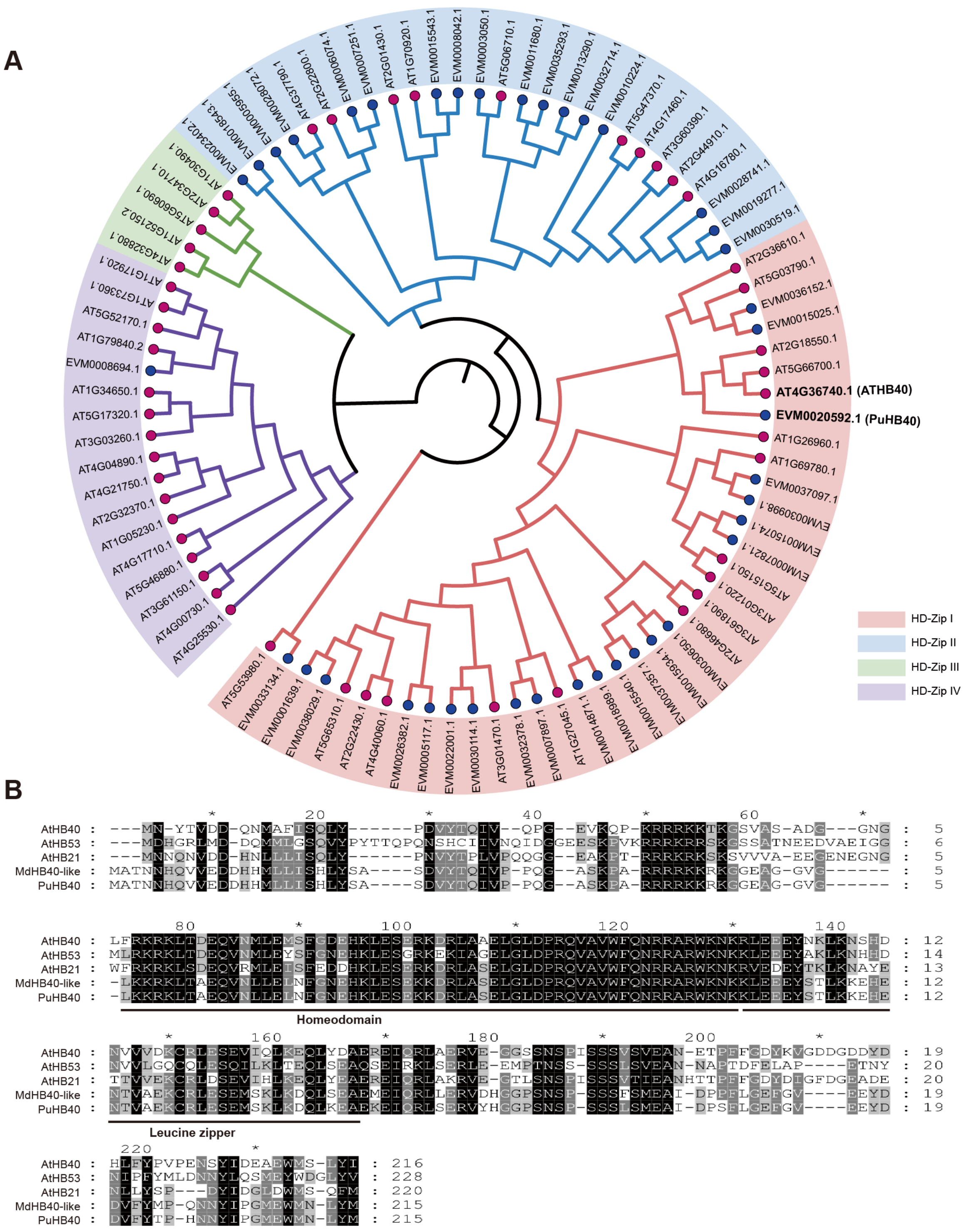


**Supplemental Figure S9. Identification of PuHB40 (Supports Figure 4). (A)** Phylogenetic analysis of candidate HD-Zip I TFs from pear and Arabidopsis. The phylogenetic tree was calculated using the Maximum-likelihood method of MEGA X (version 10.1.8). Bootstrap values of 1000 replicates for each branch are shown. Different subgroups are marked with different colors background respectively. PuHB40 (EVM0020592.1) and ATHB40 (AT4G36740.1) are bolded and enlarged. The protein sequences from pear (Pu, *Pyrus ussuriensis*, highlighted with blue dots), Arabidopsis (At, *Arabidopsis thaliana*, highlighted with rose red dots). **(B)** Amino acid sequence alignment of PuHB40 with the homologous HD-Zip I TFs from other plant species. The conserved homeodomain and Leucine zipper are shown. Related proteins including *Arabidopsis thaliana* ATHB40 (AT4G36740.1), ATHB53 (AT5G66700.1), ATHB21 (AT2G18550.1), *Malus* × *domestica* MdHB40-like (MDP0000230511), *Pyrus ussuriensis* PuHB40 (EVM0020592.1).

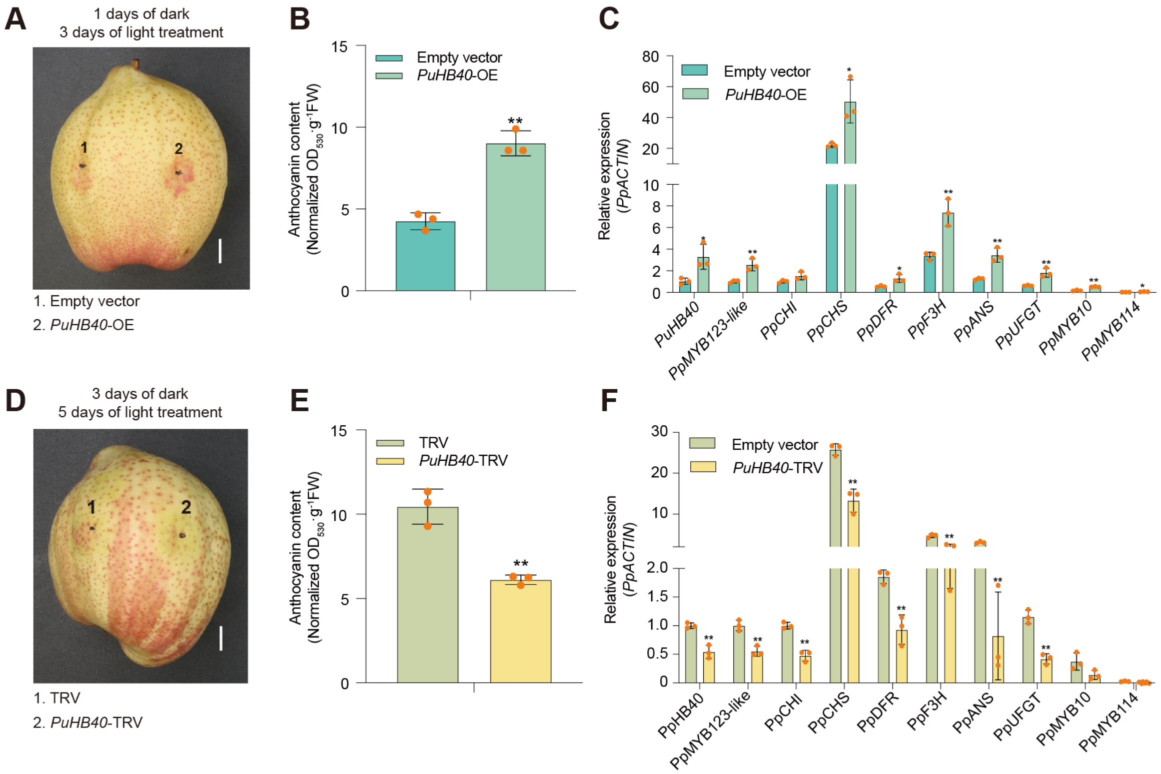


**Supplemental Figure S10**. **Functional analysis of PuHB40 in 'Hongzaosu' and 'Zaosu' pear fruits (Supports Figure 6). (A)** Transient overexpression of *PuHB40* in pear fruits (scale bar, 1 cm). The full-length CDS of *PuHB40* was inserted into the pGreenII 0029 62-SK vector under the control of the 35S promoter. Pear fruits were infiltrated with *A. tumefaciens* GV3101 cells containing the recombinant plasmid using a needle syringe. Fruits infiltrated with an empty pGreenII 0029 62-SK vector were used as the control. The phenotypes were examined after dark treatment for one day followed by the light treatment for three days. **(B)** Total anthocyanin contents in fruits transiently overexpressing *PuHB40* (units: $A_{530}$/g of fresh weight). **(C)** Expression patterns of genes related to anthocyanin biosynthesis in fruits transiently overexpressing *PuHB40*. **(D)** Transient silencing of *PuHB40* in fruits (scale bar, 1 cm). The empty vectors (pTRV1 + pTRV2) were used as the negative control. Pear fruits were placed in darkness for three days and then treated with strong light for five days. **(E)** Total anthocyanin contents in fruits transiently silencing *PuHB40* (units: $A_{530}$/g of fresh weight). **(F)** Expression patterns of genes related to anthocyanin biosynthesis in fruits in which *PuHB40* was transiently silenced. Data are presented as means ± s.d. of 3 biological replicates. Asterisks indicate significant differences compared with empty vector **(B)** and **(C)** or TRV **(E)** and **(F)** (two-tailed Student's *t-test*, $*P < 0.05$, $**P < 0.01$), all $P$ values are shown in Supplemental Data Set S7.

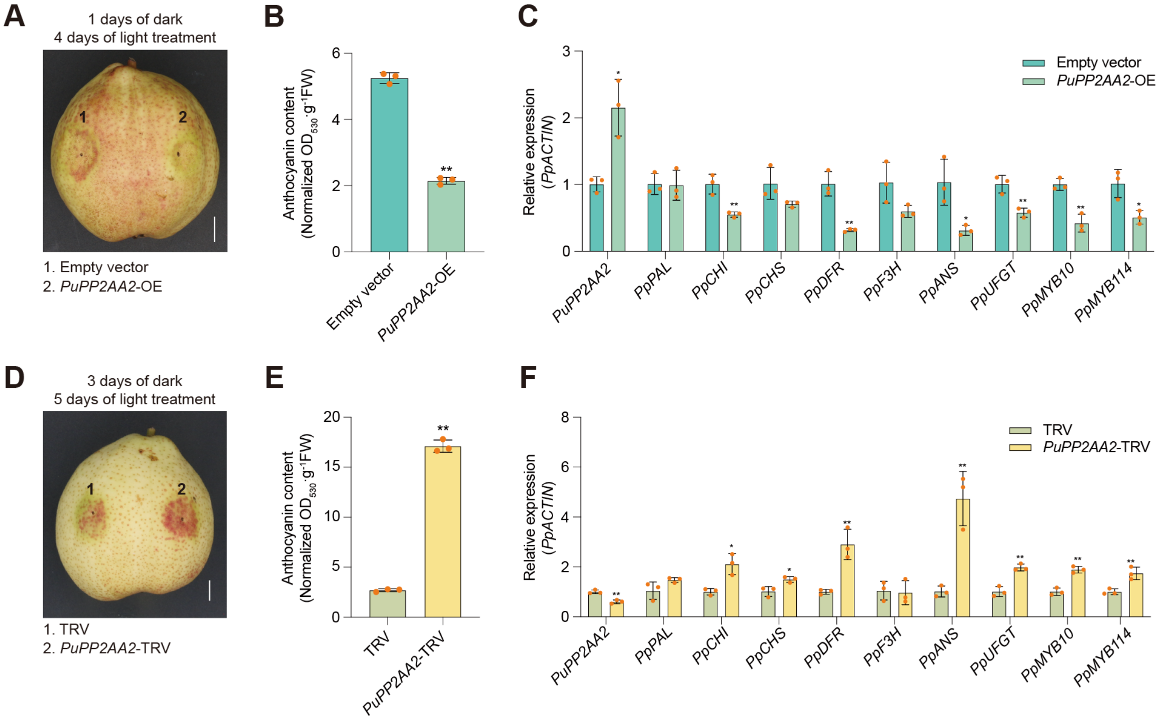


**Supplemental Figure S11. Functional analysis of PuPP2AA2 in 'Hongzaosu' pear fruits (Supports Figure 9). (A)** Transient overexpression of *PuPP2AA2* in pear fruits (scale bar, 1 cm). The full-length CDS of *PuPP2AA2* was inserted into the pGreenII 0029 62-SK vector under the control of the 35S promoter. Pear fruits were infiltrated with *A. tumefaciens* GV3101 cells containing the recombinant plasmid using a needle-free syringe. Fruits infiltrated with an empty pGreenII 0029 62-SK vector were used as the control. The phenotypes were examined after dark treatment for one day followed by the light treatment for four days. **(B)** Total anthocyanin contents in fruits transiently overexpressing *PuPP2AA2* (units: $A_{530}$/g of fresh weight). **(C)** Expression patterns of genes related to anthocyanin biosynthesis in fruits transiently overexpressing *PuPP2AA2*. **(D)** Transient silencing of *PuPP2AA2* in fruits (scale bar, 1 cm). The empty vectors (pTRV1 + pTRV2) were used as the negative control. Pear fruits were placed in darkness for three days and then treated with strong light for five days. **(E)** Total anthocyanin contents in fruits transiently silencing *PuPP2AA2* (units: $A_{530}$/g of fresh weight). **(F)** Expression patterns of genes related to anthocyanin biosynthesis in fruits in which *PuPP2AA20* was transiently silenced. Data are presented as means ± s.d. of 3 biological replicates. Asterisks indicate significant differences compared with empty vector **(B)** and **(C)** or TRV **(E)** and **(F)** (two-tailed Student's *t-test*, *$P < 0.05$, **$P < 0.01$), all *P* values are shown in Supplemental Data Set S7.
